# Complete NMR assignment for 275 of the most common dipeptides in intrinsically disordered proteins

**DOI:** 10.64898/2025.12.10.693443

**Authors:** Tobias Rindfleisch, Emilie Fjeldberg Taule, Markus S. Miettinen, Jarl Underhaug

**Author notes:** corresponding authors: Tobias Rindfleisch (,), Markus S. Miettinen, Jarl Underhaug. these authors contributed equally to this work (Markus S. Miettinen, Jarl Underhaug).

## Abstract

Accurate NMR chemical shift assignments are essential for atomic-resolution characterization of proteins. Especially for intrinsically disordered proteins (IDPs) and regions (IDRs), however, the assignment remains a labor-intensive task due to spectral overlap and conformational heterogeneity. Consequently, complete side-chain assignments are rare. Here, we present a comprehensive reference dataset, comprising the complete NMR chemical shift assignments for 275 of the most prevalent dipeptides in the IDPome, covering 93% of it. In addition, we report side-chain protonation–dependent chemical shifts for dipeptides containing aspartic or glutamic acid. The dataset contains all NMR-accessible backbone and side-chain nuclei, in total 11 571 validated data points, as well as the 1D (^1^H, ^13^C) and 2D (^1^H–^15^N HSQC, ^1^H–^13^C HSQC, TOCSY, NOESY, ^1^H–^13^C HMBC) spectra used for the assignment, making it a rich resource for the training, testing, and benchmarking of tools for data-driven protein assignment, peak picking, and synthetic spectrum generation. To facilitate such machine learning applications, all data are delivered in standardized, machine-readable formats.

## Background & Summary

Accurate chemical shift assignments lay the foundation for nuclear magnetic resonance (NMR) analysis, enabling the structural and dynamic characterization of proteins at atomic resolution. However, the assignment process remains one of the most labor-intensive steps in NMR studies, often complicated by signal overlap, peak broadening, and conformational heterogeneity^1,2^. These issues are particularly pronounced in intrinsically disordered proteins (IDPs), where the lack of a single stable tertiary structure leads to spectral congestion and poor chemical shift dispersion^3,4^. As a result, complete side-chain assignments are rarely achieved for IDPs, leaving a substantial gap in available reference data. To address this gap, a consistent and comprehensive reference set would be required, particularly one that includes all side-chain nuclei. Here, rather than relying on heterogeneous IDP datasets with incomplete assignments, we adopted a reductionist approach and focused on the smallest nontrivial building blocks of proteins—the dipeptides.

The 20 canonical amino acids, each with its unique physicochemical properties, are the elementary building blocks of proteins. Within these large macromolecules, responsible for numerous essential functions in all aspects of life, already the simple connection of just two L-amino acids via a peptide bond leads to 400 possible combinations for L-α-dipeptides. So far, these smallest nontrivial units have been poorly studied in comparison to single amino acids, longer peptides, and complete proteins^5^. However, the key characteristics of polypeptides, such as solubility and reactivity in enzymatic context, emerge from the comprising dipeptides, whose torsion angle preferences naturally also influence the protein folding tendencies. Concerning the specific case of IDPs, dipeptides are known to exhibit structural and dynamic properties similar to those of disordered chains^6^, providing an ideal minimal model for exploring disorder-related phenomena.

IDPs lack a unique and fixed tertiary (3D) structure, mainly because of the underrepresentation of hydrophobic residues, which prevents the formation of a hydrophobic core, coupled with the enrichment of charged and polar amino acids, which promotes protein–solvent interactions^7,8^. However, their broad structural ensembles do possess characteristic sequence-dependent distributions over the conformational space^4^. Not all IDPs are fully unstructured; the term also includes proteins with intrinsically disordered regions (IDRs) embedded in otherwise well-folded domains^9^. Bioinformatic analyses predict more than 40% of eukaryotic proteins to contain IDRs^10^ with approximately two thirds of these being fully disordered^11^. Achieving atomistic-resolution characterization of IDP ensembles remains a major challenge, as their continuous rapid interconversion between innumerable conformations prevents the use of traditional structure-determination techniques such as X-ray crystallography or cryo-electron microscopy^8,12–14^.

Among experimental techniques, NMR spectroscopy uniquely enables atomistic interrogation of IDP ensembles^15^, especially when coupled with all-atom molecular dynamics (MD) simulations^16–20^. The development and validation of MD models (force fields), as well as a broad range of other computational protein-modeling approaches, rely on rigorous comparison against experimental observables, making chemical shifts a valuable resource in the structural context^16–20^. Such validational use, however, requires precise and reliable NMR assignments, in which the observed chemical shifts are unambiguously matched to their corresponding atoms. Already backbone shifts provide a useful benchmark, but their limited number restricts the depth of model evaluation. Incorporating side-chain chemical shifts would greatly enhance the statistical power for testing and refining theoretical models; yet this potential remains largely unrealized due to the mentioned side-chain assignment gap in IDPs. Previous studies on mono-^21^, di-^6^, tri-^22^, and small oligopeptides^23^ have shown most of these molecules to reflect the behavior of IDPs, suggesting dipeptides as the simplest unit that can capture the fundamental sequence-dependent correlations governing the intrinsic disorder.

Here, we provide complete chemical shift assignments for 275 of the most frequently occurring dipeptides in the IDP proteome (IDPome), covering 93% of it. The dataset, comprising 11 571 unique data points quality-controlled by manually cross-checking multiple 2D-NMR spectra, includes all NMR-accessible nuclei, in both the backbone and the side-chains, thereby bridging the IDP-side-chain data gap. We further assessed side-chain protonation effects on chemical shifts by measuring dipeptides containing aspartic or glutamic acid with both protonated and deprotonated side-chain carboxyl groups. Crucially, we also provide the complete set of acquired and processed spectra—1D (^1^H and ^13^C) as well as 2D (^1^H–^15^N HSQC, ^1^H–^13^C HSQC, TOCSY, NOESY, and ^1^H–^13^C HMBC)—used for assignment. These enable the development and benchmarking of algorithms for automated all-atom assignment of IDPs and the generation of synthetic NMR spectra for (partial) protein sequences. The dataset also provides a benchmark for improving automated peak-picking methods, including the deconvolution of overlapping resonances. By delivering standardized, high-quality, machine-readable data, this resource advances reproducibility in both computational and experimental NMR studies, facilitating progress across the protein disorder spectrum.

## Methods

### Sample selection

To ensure a comprehensive representation of IDPs and efficient use of resources, we aimed to include the most frequently occurring dipeptides in the IDP proteome. This strategy inherently excludes the most hydrophobic dipeptides, minimizing potential solubility issues and allowing all samples to be measured under standardized experimental conditions using a single, uniform solvent.

To find the most frequently occurring dipeptides, we evaluated the occurrence of all 400 canonical dipeptides in the IDPome by analyzing the complete DisProt^24^ database (disprot.org, release version 2025-06) that exclusively contains experimentally confirmed IDP sequences. For each dipeptide, we counted the absolute occurrence across all non-redundant sequences and determined the corresponding relative frequencies. As DisProt includes redundant entries, derived from multiple experimental reads, under the same protein-sequence identifier (ID), only the first sequence associated with each ID was considered (scripts are provided; see section Code Availability). We then ranked the 400 dipeptides according to descending relative frequency.

The order in which we then performed our experiments followed the frequency ranking we had obtained—to the extent that was feasible in practice—but all proline-containing dipeptides were included due to the distinctive conformational characteristics of proline in IDPs^25,26^.

### Sample preparation

Unlabeled dipeptides with N-terminal acetylations and C-terminal amidations were purchased from Peptide Protein Research Ltd, Fareham, United Kingdom. The capping groups were added to eliminate the terminal charges and to introduce a third peptide bond, as well as a third C^*α*^, thereby mimicking the chemical context within a polypeptide chain. All lyophilized samples were dissolved unbuffered in 90% H_2_O, 10% D_2_O, and 15 µM DSS (sodium trimethylsilylpropanesulfonate) at concentrations of approximately 5 mg/mL. To adjust the carboxyl group protonation states of the ASP and GLU side chains, the sample pH was changed by NaOH or HCl titration to 2.6–3.4 (protonated) or 5.7– 6.5 (deprotonated) and verified using a pH meter. The measurements were performed in 5-mm Bruker (Billerica, MA, United States) NMR tubes with a volume of 600 µL per sample.

### NMR experiments

High-throughput NMR measurements for dipeptide assignments were performed on two Bruker BioSpin spectrometers (Table S1): an Ascend 600 MHz with a QCI-P CryoProbe (219 samples) and an Ascend 850 MHz with a TCI CryoProbe (56 samples), both equipped with an AVANCE NEO console. For samples measured on the 850 MHz instrument, additional 1D-^1^H (zgesgppe) spectra were acquired at 600 MHz. Samples were handled automatically with a SampleJet sample changer and equilibrated to 298.0 K for at least 2 minutes prior to measurement and otherwise stored at 4°C. Whenever non-uniform sampling (NUS) was used, the sampling schedule was randomly created by TopSpin’s NUS generator (seed 54321). Table 1 lists the performed NMR experiments, along with their key acquisition parameters.

**Table 1.**
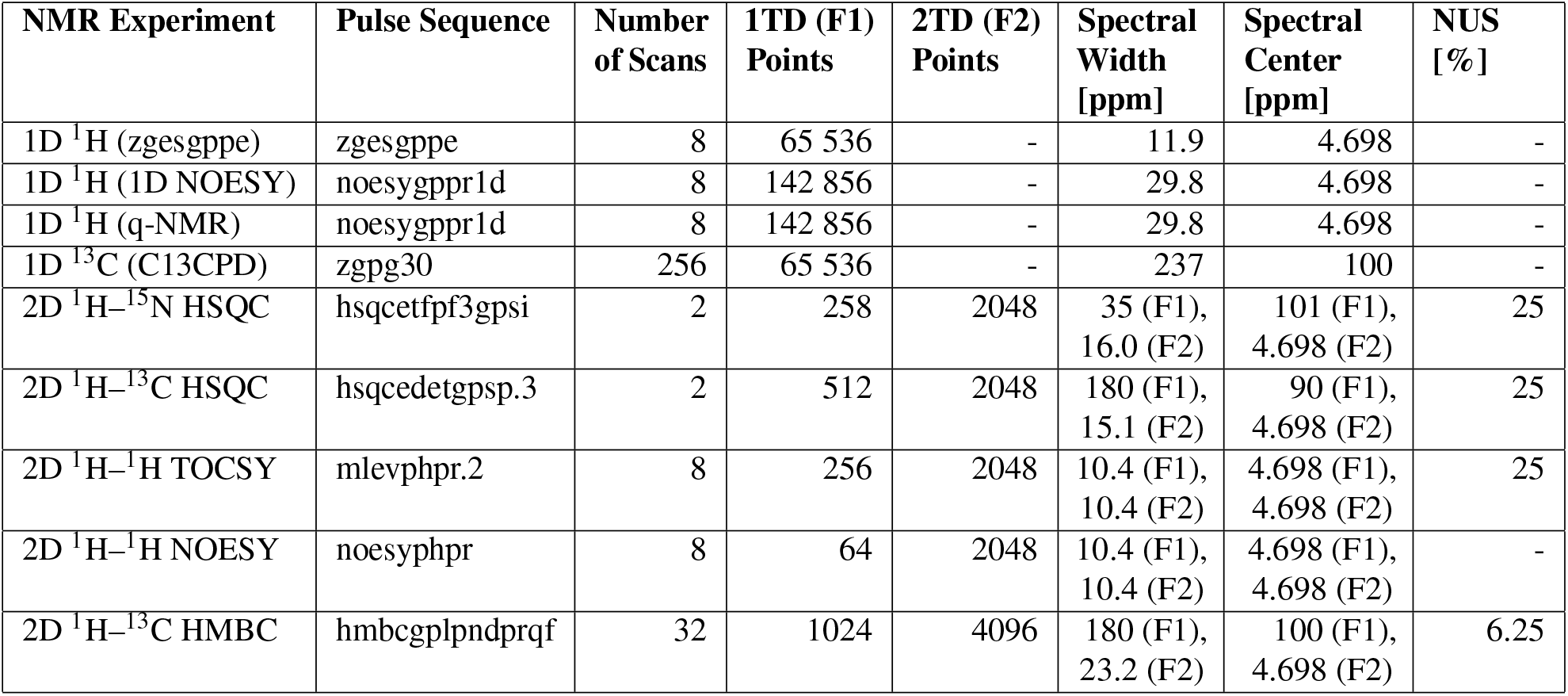
Overview of selected acquisition parameters for dipeptide NMR assignment experiments: number of scans (NS), number of acquisition points in the F1 and F2 dimensions (1TD and 2TD), spectral width (SW), spectral center (O1P), and the applied percentage of non-uniform sampling (NUS). All pulse sequences were included in TopSpin (version 4.1.4, Bruker BioSpin). For the complete list of acquisition and processing parameters per sample, see the Edmond repository^27^.

### Spectra processing and assignments

Acquired NMR spectra were processed using the MestReNova software (version 14.02.0-26256, Mestrelab Research) and referenced to DSS. NUS-acquired spectra were reconstructed using MestReNova’s MIST algorithm (Modified IST, based on Interactive Soft Thresholding (IST)^28^) in static mode. Bruker datasets (Bruker BioSpin), containing the free induction decays (FIDs), were imported into MestReNova and processed with standard routines; the processing steps included zero-filling (doubling the number of data points in 1D spectra and in the indirect dimension of 2D spectra), application of window functions, Fourier transformation, phase correction, and baseline correction. For 1D spectra, an exponential multiplication (EM) window function was applied with line broadenings of 0.5 Hz (zgesgppe) or 0.3 Hz (1D NOESY and q-NMR) for ^1^H, and 1 Hz for ^13^C. For 2D spectra, a squared cosine function was used. All spectra were baseline corrected using a third-order polynomial fit, except for the 1D ^13^C (C13CPD) experiments, where a Whittaker smoother was applied. In 2D experiments, the residual water resonance was suppressed using MestReNova’s signal suppression tool, selecting the solvent (water) frequency and applying a narrow suppression filter to remove the water-signal interference.

Chemical shifts of dipeptides were manually assigned for all detectable ^1^H, ^13^C, and ^15^N nuclei using established sequential assignment strategies. Precise values for ^1^H and ^13^C chemical shifts were derived from the corresponding 1D spectra—for non-separable, overlapping multiplets ^1^H–^13^C HSQC spectra were considered—and ^15^N shifts were obtained from the F1 (^15^N) dimension of ^1^H–^15^N HSQC experiments. Clearly identifiable multiplets were analyzed with MestReNova’s multiplet manager, and final assignments were validated by cross-checking correlations among several spectra.

NMR spectra with respective acquisition and processing parameters were stored in the machine-readable NMReData^29^ format, and the corresponding peak lists were exported as Structure Data Files (SDF) and converted to the NMR-STAR^30^, YAML, and chemical shift table formats, all compliant with the Biological Magnetic Resonance Bank (BMRB, bmrb.io) nuclei naming standards^31^.

## Data Records

The chemical shift datasets for 275 of the most common dipeptides in IDPs, along with the corresponding NMR spectra (Table 1), are permanently openly available in the Edmond Open Research Data Repository (edmond.mpg.de) entry DOI: 10.17617/3.CBC-NQG^27^. The top level of the deposition contains a `Data` directory storing all assignment related files, along with a second `Script` directory including a fully automated Snakemake^32^ workflow that executes the complete data-processing, analysis, and visualization pipeline. In addition, the `Script` directory contains three subdirectories that include the individual scripts used for (i) dipeptide ranking, (ii) creation of the dipeptide dataset files, and (iii) generation of all figures presented in this publication.

The `Data` directory includes 20 subdirectories named after the canonical amino acids, each representing the first residue of the dipeptides, e.g., `Data/A/`. Within each of the 20 subdirectories, the corresponding dipeptide datasets are stored in their own subdirectories: `Data/A/AA/`. Each dipeptide dataset includes the acquired, processed, and analysis-related files in machine-readable formats (see Table 2), facilitating reuse of the NMR data. For ASP- and GLU-containing dipeptides, two subdirectories are provided, each storing datasets for samples with either protonated or deprotonated side-chain carboxyl groups: `Data/A/AD_protonated/` and `Data/A/AD_deprotonated/`. For samples measured on the 850 MHz instrument (Table S1), the corresponding datasets also contain a `600MHz` directory, which stores a single 1D-^1^H (zgesgppe) experiment (`.mnova` and `_nmredata.zip` files) acquired at 600 MHz, together with the corresponding ^1^H chemical shift assignments (`_Chemical_Shifts.txt` and `_Chemical_Shifts.yml` files).

**Table 2.**
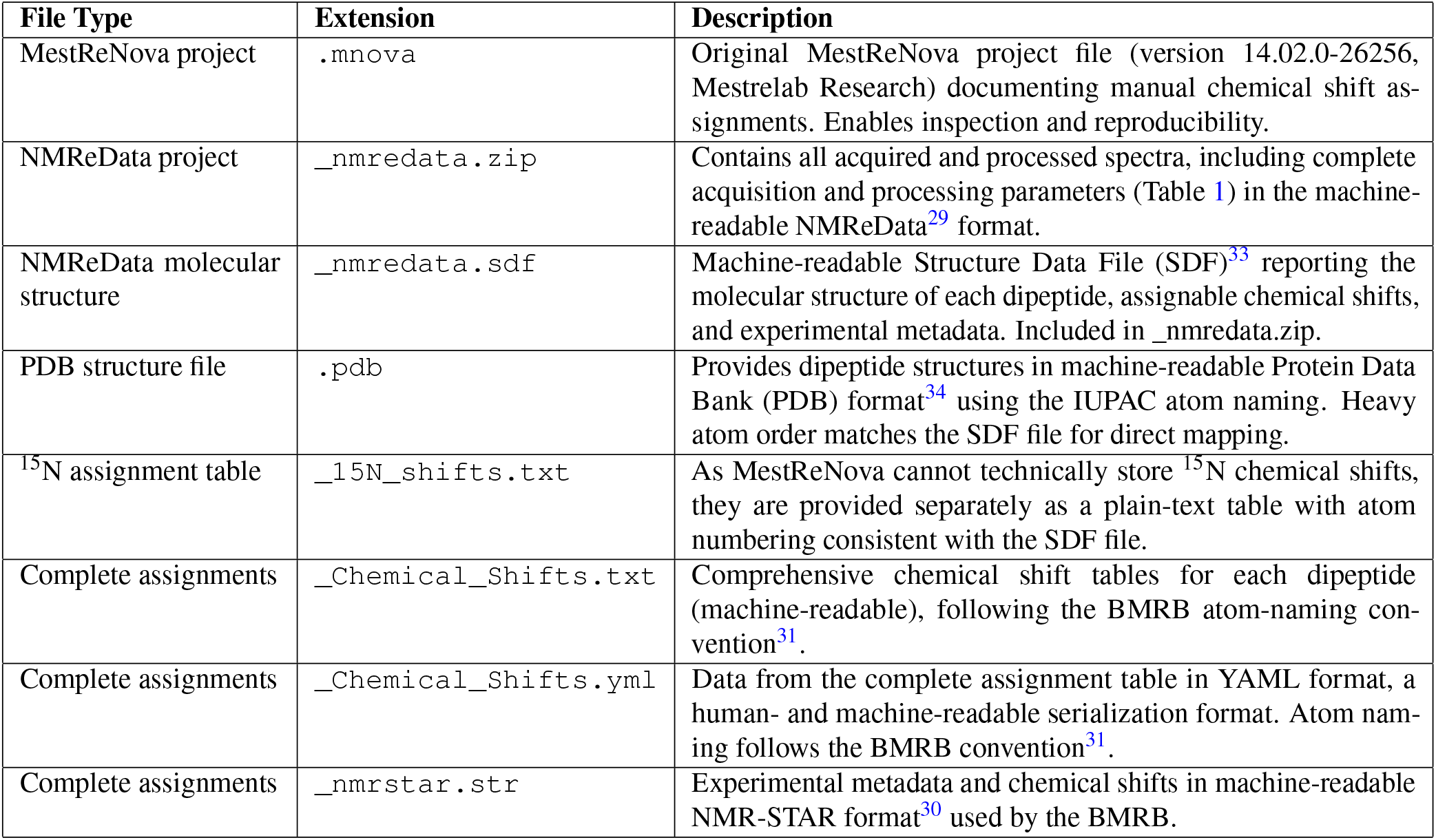
Overview of the deposited dipeptide dataset files and their contents.

Since the assignment of diastereotopic methylene protons is inherently ambiguous, we followed the common practice of reporting them at the CH_2_ group level as HX’ and HX” (values are interchangeable, X is a placeholder for any side-chain position) in the `_Chemical_Shifts.txt` and `_Chemical_Shifts.yml` files (Table 2). In the NMR-STAR format (`_nmrstar.str` files, Table 2), the interchangeable character of both protons is reflected by the BMRB Ambiguity Index^31^ of “2” (geminal atom ambiguity) for the corresponding HX2/HX3 notation. For consistency across files and figures, we list the lower-frequency proton as HX’/HX2 and the higher-frequency proton as HX”/HX3. In figures, the diastereotopic ^1^H shifts remain grouped under the label Q, with both protons shown separately to avoid misinterpretation due to their interchangeable character.

## Data Overview

The frequencies of all 400 canonical dipeptides in IDPs are visualized in Figure 1 and listed in Table S2. These frequencies were derived from an analysis of 3 201 non-redundant and experimentally confirmed IDP sequences included in the DisProt database^24^ (release version 2025-06). These contained 221 637 individual dipeptides, providing a quantitative overview of dipeptide occurrences across the IDP proteome. According to their cumulative relative frequency (Figures 1, S1, and S2), the 275 dipeptides we study here collectively account for 93% of the IDPome, faithfully reflecting the characteristic amino acid composition and physicochemical properties of disordered proteins and regions.

**Figure 1.**
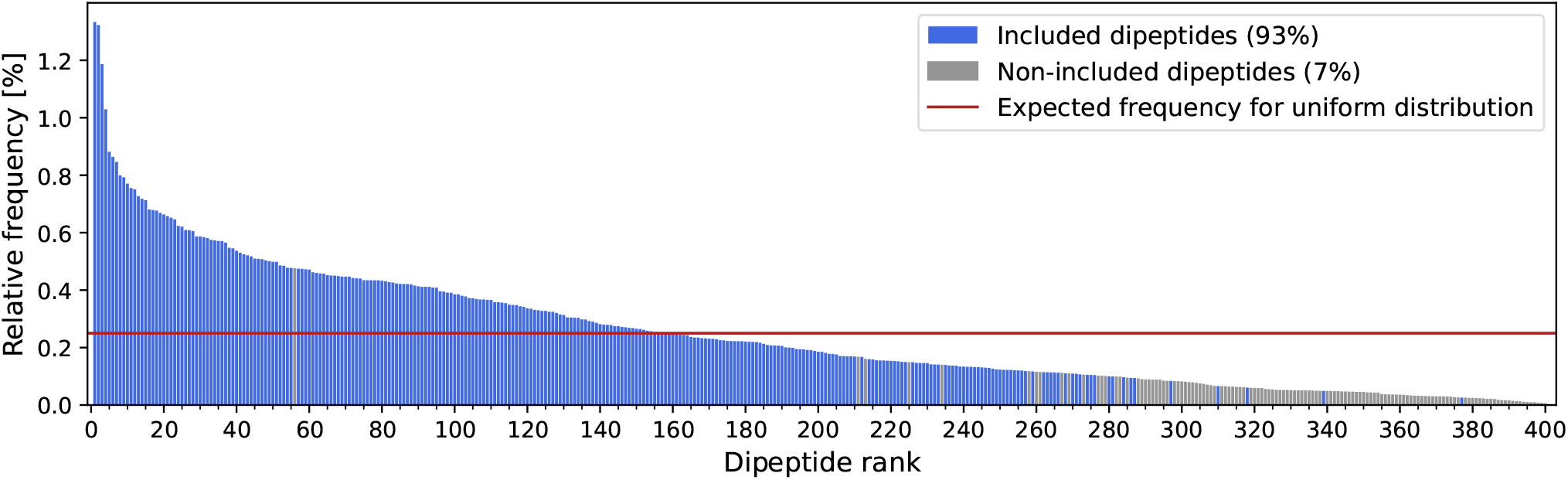
Relative frequency distribution of the 400 canonical dipeptides in the IDPome, as derived from the DisProt^24^ database, compared to the uniform distribution (red line). Blue bars indicate dipeptides included in the presented dataset, gray the non-included ones; total percentages of both populations are shown in the legend keys. Figure S1 shows the same plot with the explicit dipeptide names instead of the rank on the *x*-axis. Figure S2 shows the cumulative relative frequency as a function of the dipeptide rank. Table S2 connects the dipeptide rank with the dipeptide names and additional details, including relative frequencies and absolute occurrences.

Dipeptide frequencies are non-uniformly distributed, with only a small subset occurring at the 0.25% average occurrence; approximately one third of the dipeptides appear more frequently than this, while about half occur more seldomly (Figure S1). The 86 most common dipeptides—roughly one fifth of all the 400—account for half of the entire IDPome (Figure S2), underscoring the compositional bias in disordered proteins. As expected, the most frequently observed dipeptides contain small and/or non-hydrophobic residues (Figure S1 and Table S2), whereas those incorporating one large hydrophobic residue are less common and those containing two are rare. Cysteine- and tryptophan-containing dipeptides are among the least represented, consistent with their structure-stabilizing roles that are incompatible with the disordered nature of IDPs^35^. Consequently, although roughly 100 dipeptides—one quarter of all possible combinations—contain large hydrophobic residues or cysteine, they collectively account for less than 7% of the IDPome, resulting in the high coverage of the here presented dataset.

Note, however, that a few high-ranking dipeptides were left out of the dataset, because the samples either (i) contained substantial impurities (AT) preventing reliable chemical shift assignments, (ii) were insoluble (FL, IV, IN) under the chosen solvent conditions, or (iii) exhibited extensive peak overlap (II).

## Technical Validation

All chemical shifts were assigned manually and systematically verified across multiple spectra. Any ambiguous cases were discussed and resolved by at least two independent researchers to minimize subjective bias. To ensure completeness and consistency, a verification script was used to examine the assignment tables and NMR-STAR files for missing or malformed entries, confirming that all the expected NMR-active nuclei were present (scripts are provided^27^; details in section Code Availability). The measurement uncertainty of chemical shifts under constant experimental conditions is typically minimal to negligible^36–38^, and referencing to DSS ensures that systematic errors are effectively eliminated. Cross-validation was achieved using a comprehensive set of 2D experiments (Table 1), with TOCSY connectivity ensuring that assignments between identical residues in homodipeptides could not be inadvertently swapped.

We further validated the assigned chemical shifts by direct comparison to the BMRB’s amino-acid-wise statistics^31^ (bmrb.io, accessed October 2025), which include a BMRB-wide average for every detectable NMR-active nuclei: For each assignable nucleus across the 275 dipeptides, we calculated the difference between the observed chemical shift (*δ*_obs_) and the corresponding BMRB average shift (reference; *δ*_ref_) as Δ*δ* = *δ*_obs_ −*δ*_ref_. Figure 2 shows the individual nuclei-distributions of Δ*δ* (points) for each of the 20 amino acid residues; the shaded intervals mark the standard deviation (SD) of the corresponding BMRB reference distribution as ±1SD (light gray) and ±2SD (dark gray). It is seen that 95.6% of our observed data points are within ±1SD of the BMRB reference (Figure 2), and all but one within ±2SD; the deviating backbone ^15^N chemical shift of deprotonated ASP in the dipeptide DP was confirmed in two independent samples and may result from the combination of two factors: the negative ASP side-chain charge in close proximity to the PRO backbone. Overall, the comparison to BMRB reference shifts indicates a minimal likelihood of misassignments, which is further supported by the absence of major outliers (Figure 2); only a few, reliably explainable exceptions are present.

**Figure 2.**
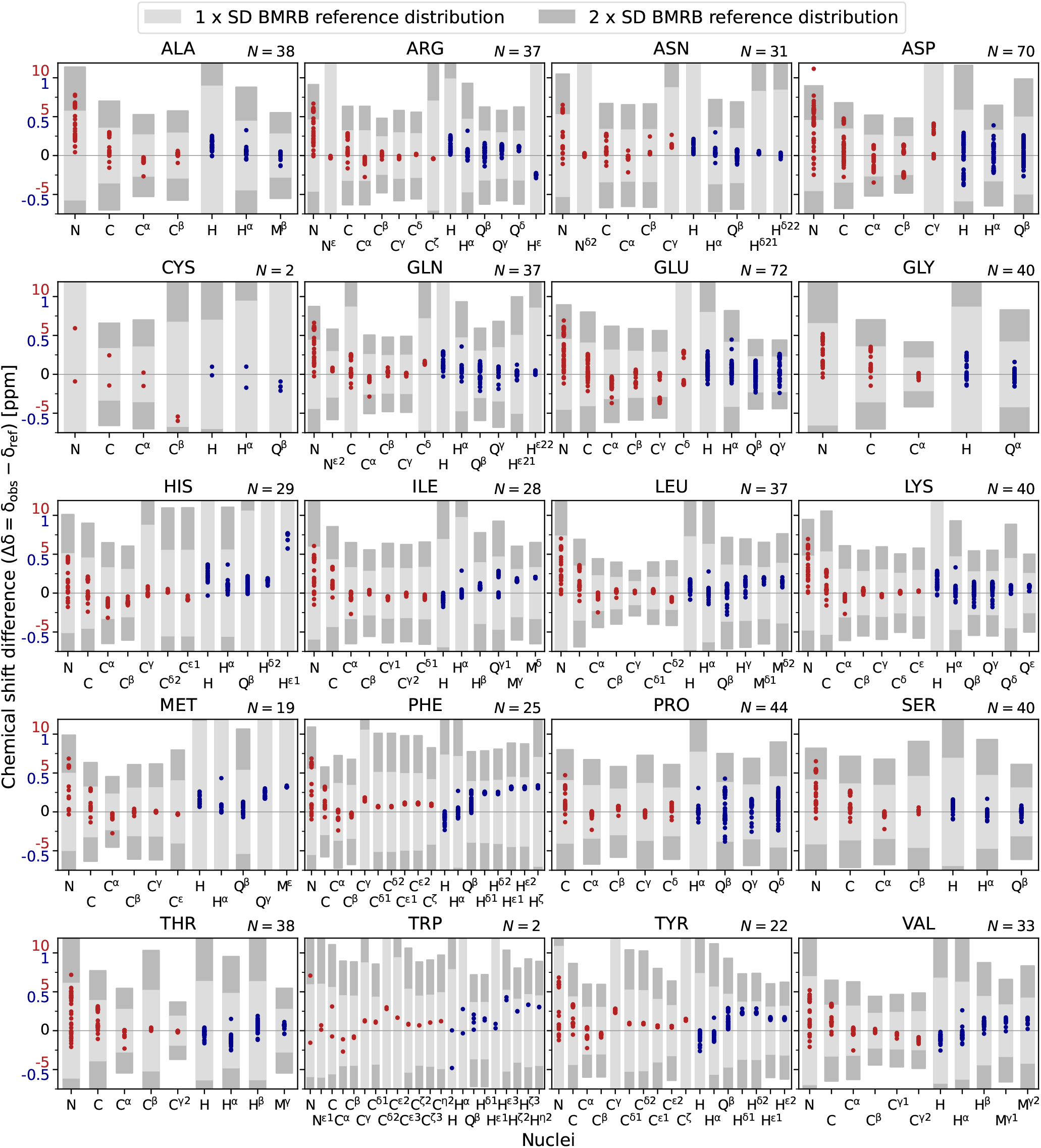
Comparison to BMRB: The 20 residue-specific panels show Δ*δ* = *δ*_obs_ − *δ*_ref_ for each assignable nucleus, defined as the difference between the observed chemical shifts (*δ*_obs_) and the corresponding BMRB reference values^31^ (*δ*_ref_). Each panel has two *y*-axis scales, since the ^1^H Δ*δ* (blue data points and labels) span a much narrower range than the ^13^C and ^15^N ± Δ*δ* (red). Shaded regions show the standard deviation (SD) intervals of the respective BMRB reference distributions^31^: ± 1SD (light gray) and ±2SD (dark gray). *N* is the total number of observations (one/two per measured hetero/homodipeptide) per residue; dipeptides with protonated and deprotonated ASP or GLU side chains are considered separately (Figure S3 summarizes the influence of side-chain protonation). ^1^H resonances are classified as H, Q, and M: H represents the CH protons with one data point; Q the CH_2_ groups, for which the two protons are shown as separate data points; and M the methyl protons, which generally yield a single averaged chemical shift.

Note that in the ^1^H–^15^N HSQC experiments, the ^15^N spectral range did not cover the chemical shifts of arginine N^*ε*^ nuclei (signal at roughly 85 ppm), leading this resonance to appear near 120 ppm due to aliasing (folding)^39^; we corrected this by subtracting the spectral width (35 ppm) from the observed resonances. As this correction is frequently omitted in the BMRB, biasing the averages reported there toward higher values, we used 85 ppm as the reference value for arginine N^*ε*^ nuclei in our technical validation (Figure 2). Another common issue in BMRB reference data concerns the side-chain amide groups of asparagine (H^*δ*21^ and H^*δ*22^) and glutamine (H^*ε*21^ and H^*ε*22^), where the two protons are often swapped in assignments. To avoid this ambiguity, we used the centers of the BMRB-reported H^*δ*21^/H^*δ*22^ and H^*ε*21^/H^*ε*22^ populations as the reference values.

In Figure 2, the backbone ^15^N nuclei exhibit the largest systematic deviations, reflecting their dependence on the preceding residue^40^. A similar but less pronounced dependency is observed for backbone carbonyl ^13^C chemical shifts^40^: While some ^15^N backbone shifts are not within the ±1SD interval, this only occurs marginally for the ^13^C carbonyls (C) of ASP, LEU, PHE, and VAL.

Overall, side-chain protonation of ASP and GLU primarily induces intra-residue effects (Table S3) and only minor changes in neighboring residues (Table S4), although these still exceed typical chemical shift assignment uncertainties. Pronounced intra-residue effects are seen for the ASP backbone and side chain (Table S3), with deprotonated states exhibiting larger ^13^C and ^15^N chemical shifts and reduced ^1^H shifts (Figure S3). GLU shows similar protonation-dependent behavior, but the effects remain largely limited to the side chain and couple only weakly to the backbone.

Closer examination reveals consistent sequence-dependent trends. Across all residue-specific plots of Figure 2, H^*α*^ and C^*α*^ have a single data point deviating from the main cluster(s). All these arise from dipeptides with proline in the second position (X–PRO-motif, with X being any amino acid; Figures S4 and S5). This systematic behavior reflects the altered chemical environment introduced by proline, which lacks an amide proton and modifies the N–C connectivity within the peptide bond.

The main clusters of carbonyl C, backbone amide N, and C^*α*^ chemical shifts separate into two distinct populations (Figures S6 and S7), corresponding to the residue’s position within the dipeptide (N-terminal versus C-terminal). The absence of such splitting beyond C^*β*^ in side-chains is consistent with the expectation that residue-position effects should primarily affect the backbone nuclei.

Distinct ^1^H resonance populations are consistently observed near aromatic residues (PHE, TRP, and TYR), reflecting the influence of a local environment with an increased electron density. This effect affects all nuclei of residues connected to an aromatic residue and is most pronounced for H^*β*^ and H^*γ*^ nuclei, showing decreased ^1^H chemical shifts. Smaller but still pronounced effects are observed on backbone H, H^*α*^, H^*δ*^, and H^*ε*^ nuclei (Figures S6 and S7).

For some nuclei (e.g., in PHE or TYR), certain chemical shift clusters show more distinct deviations from the BMRB-wide average than others (Figure 2), which is largely attributed to their non-Gaussian nature: Since BMRB distributions are sums of several overlaying populations, corresponding to different protein, solvent, or measurement properties, their global mean does not necessarily represent the centers of the underlying sub-distributions. Consequently, dipeptide chemical shifts that populate regions farther from the global BMRB average appear deviating, yet being fully consistent within their population. To address this heterogeneity in BMRB distributions, we performed our technical validation not only based on the deviation from the BMRB average, but also included its ±1SD (light gray) and ±2SD (dark gray) intervals (Figures 2 and S3–S7).

Figure S8 indicates that variations in chemical shifts between measurements at pH ≈3 and ≈6 are small for those residues whose protonation states remain unchanged. As these measurements were performed only on dipeptides containing ASP or GLU, the observed differences may reflect both pH effects and changes in ASP/GLU protonation. However, because the observed chemical shift variations differ in magnitude between ASP and GLU, with more pronounced perturbations for ASP-containing dipeptides, they appear to be primarily driven by ASP/GLU protonation rather than by pH. This agrees with the fact that the non-buffered sample preparation—which enables straightforward use of these data as benchmarks for molecular modelling, but may introduce some pH variability—does not introduce marked variation in the chemical shift values (Figure S3).

Changes in side-chain protonation are expected to substantially alter chemical shifts, as observed for ASP and GLU (Table S3). In contrast, no comparable changes are detected for other residues in ASP- and GLU-containing dipeptides measured between pH ≈3 and ≈6, indicating that their protonation states remain unchanged across this range (Table S4 and Figure S8). Further, their chemical shifts agree with those observed in dipeptides lacking ASP or GLU (Figure S3), supporting the conclusion that these (uncharged as well as LYS and ARG) residues adopt their fully protonated state throughout all measurements. An exception could be expected for HIS, whose pK_a_ (≈6.05) lies at the upper end of the investigated pH range; also for HIS, however, changes comparable to those in ASP and GLU are not observed (Figure S8). Furthermore, no major divergence in BMRB-difference distribution widths are observed between charged and non-charged residues across the ^1^H, ^13^C, and ^15^N shifts (Figure 2). Overall, the BMRB-difference distributions are narrow, particularly when residue-position effects are taken into account (Figures S6 and S7).

Use of NUS in 2D NMR can introduce spectral artifacts, particularly when the sampling density is insufficient. We verified that the NUS fractions applied in our ^1^H–^15^N HSQC, ^1^H–^13^C HSQC, and ^1^H–^13^C HMBC experiments only led to minor artifacts with low occurrence and did not cause any chemical shift perturbations relative to uniform sampling (Figure S9). Moreover, any influence of NUS on the ^1^H and ^13^C chemical shifts can be generally excluded, since the corresponding values were assigned from NUS-free 1D spectra.

We ensured technical transferability of chemical shift assignments between the two instruments by DSS referencing and careful temperature calibration, thereby minimizing potential instrument- or field-dependent effects, including those affecting methylene protons. To verify this, we acquired 1D-^1^H spectra at 600 MHz for all the 56 samples originally measured at 850 MHz and reassigned the ^1^H chemical shifts. The average differences between both fields were 0.0017±0.0024 ppm for all protons (n = 1 035), 0.0024±0.0024 ppm for methylene protons (n = 346), and 0.0013±0.0024 ppm for non-methylene protons (n = 689). Since the values are well within the expected ^1^H-assignment uncertainties (±0.001 to 0.01±ppm), and their error bars overlap, no significant instrument- or field-dependent effects on the ^1^H shifts are evident.

In summary, these validations corroborate the technical quality and reliability of the reported chemical shift dataset.

## Usage Notes

The presented dataset delivers complete and standardized chemical shift assignments for 275 of the most common dipeptides found in IDPs, establishing a unique reference resource comprising 11 571 rigorously quality-controlled data points. As the corresponding 1D and 2D spectra are included, it further supports robust training and validation of artificial intelligence (AI) and machine learning approaches in protein NMR spectroscopy.

In contrast to most protein NMR datasets, which typically report only backbone resonances together with C^*β*^ /H^*β*^ nuclei, our dataset provides complete assignments for all NMR-accessible nuclei in each dipeptide. To our knowledge, this represents the first protein-related dataset that achieves truly all-atom assignments, thereby bridging the side-chain data gap. Notably, the chemical shift distributions of longer-side-chain atoms (Figure S10) are remarkably narrow, reflecting generic sequence independence of these chemical shifts; nevertheless, the distributions are wider than the assignment uncertainties, which supports the feasibility of extending data-driven all-atom assignment and prediction strategies to disordered proteins. In addition, the dataset can serve as a valuable embedding or validation reference for existing NMR-related cross-validation tools (such as PANAV^41^), automated assignment software (ARTINA^42,43^), primary-sequence-based approaches for IDP random coil chemical shift prediction (ncIDP^44^ and POTENCI^45^), as well as programs connecting chemical shifts and protein structure (TALOS+^46^/TALOS-N^47^, SPARTA+^48^, SHIFTX2^49^, and PPM^50^).

In addition to the chemical shift assignments, the dataset includes the complete set of acquired and processed spectra for all the 275 dipeptides. For each sample, eight spectra are provided, covering 1D ^1^H, 1D ^13^C, and a diverse range of 2D experiments, including ^1^H–^15^N HSQC, ^1^H–^13^C HSQC, TOCSY, NOESY, and ^1^H–^13^C HMBC. This comprehensive spectral coverage greatly expands the scope of potential applications: The data can contribute to training of models to perform atom-specific assignments from experimental spectra, but also to generate synthetic NMR spectra for (partial) protein sequences, encompassing all detectable nuclei. Beyond these applications, the dataset provides a benchmark resource for advancing peak-picking algorithms, with particular potential to improve the deconvolution of overlapping peaks—a long-standing challenge in protein NMR spectroscopy^1,2^. To this end, the dataset holds potential to be integrated into ongoing NMR initiatives aiming to address these challenges, such as NMRFAM-SPARKY^51^ and the CCPN framework^52^ within the CCPN project^53^.

We expect the dataset to be especially valuable for advancing AI methods tailored to IDPs, a class of proteins whose conformational heterogeneity poses significant challenges for traditional NMR approaches. The dataset also provides a rigorous benchmark for comparative studies, enabling systematic evaluation of existing computational NMR methods in terms of accuracy, completeness, and robustness. Its standardized nature ensures reproducibility and facilitates integration with other structural biology resources—such as BMRB^31^ due to the NMR-STAR-format used^30^.

Files reporting chemical shifts (`_Chemical_Shifts.txt`, `_Chemical_Shifts.yml`, and `_nmrstar.str`; see Table 2) are provided in widely used formats and can be inspected with any standard text editor. For programmatic access, `_Chemical_Shifts.txt` files can be imported in Python using the `pandas` package, `_Chemical_Shifts.yml` files with `PyYAML`, and `_nmrstar.str` files with `pynmrstar`. All acquired and processed spectra, together with the full acquisition and processing parameters (Table 1), are stored in NMReData format (`_nmredata.zip`). This standardized format facilitates interoperability and can be opened in several software environments, including MestReNova (Mestrelab Research), TopSpin (Bruker BioSpin), and the NMReData project tools^29^ ([NMReData Initiative](https://nmredatainitiative.github.io/Main_Page/)). In addition, programmatic access and downstream processing can be carried out in Python using the `nmrglue` package^54^. Examples for programmatic access are given in the provided scripts (see section Code availability).

## Data availability

The dataset containing the chemical shifts for 275 of the most common dipeptides in IDPs, along with the corresponding NMR spectra used for their assignment, is permanently and openly available on the Edmond Open Research Data Repository (edmond.mpg.de) entry^27^ DOI: 10.17617/3.CBCNQG.

## Code availability

The code of the complete computational pipeline—including (i) the analysis of dipeptide occurrence frequencies in DisProt, (ii) the generation of NMR chemical shift report files (`_Chemical_Shifts.txt`, `_Chemical_Shifts.yml`, and `_nmrstar.str`; see Table 2), and (iii) the creation of all figures—is permanently and publicly available on the Edmond Open Research Data Repository (edmond.mpg.de) entry^27^ DOI: 10.17617/3.CBCNQG.

The implementation within a reproducible Snakemake^32^ workflow ensures transparent and automated execution of all analysis and visualization steps. The code used to generate Figure 2 is additionally provided as an example template for accessing the dataset.

Versions 4.3, 4.4, and 4.5 of TopSpin (Bruker BioSpin) and version 14.02.0-26256 of MestReNova (Mestrelab Research) were used for data acquisition and processing. Python 3.9.13 was employed to generate all figures and files reporting chemical shifts. The following Python packages were used: `biopython` (1.78), `matplotlib` (3.5.2), `numpy` (1.21.5), `os` (3.9.13), `pandas` (1.4.4), `pynmrstar` (3.3.5), `pyYAML` (6.0), `re` (3.9.13), `tqdm` (4.64.1), and `zipfile` (3.9.13). These details are also included in the headers of the provided scripts.

## Acknowledgements

We thank Prof. Dr. Reinhard Lipowsky and Dr. Hanne Antila (Theory and Biosystems, Max Planck Institute of Colloids and Interfaces) for invaluable support in purchasing the dipeptide samples. We thank Prof. Dr. Dirk Walther (Max Planck Institute of Molecular Plant Physiology) for support and valuable discussions.

## Author contributions statement

J.U., T.R., and M.S.M. conceived the NMR experiments; T.R. performed computational analysis for sample selection; M.S.M provided the samples; T.R., E.F.T., and J.U. conducted the NMR experiments; T.R., E.F.T., and J.U. analyzed the NMR results; T.R. and J.U. performed technical validation and deposited the NMR data; J.U. and M.S.M. supervised the research; T.R. wrote and M.S.M edited the manuscript with inputs from all co-authors. All authors reviewed the final manuscript.

## References

1. Bax, A. & Grzesiek, S. Methodological advances in protein NMR. Accounts Chem. Res. 26, 131–138 (1993).

2. Dyson, H. J. & Wright, P. E. Unfolded proteins and protein folding studied by NMR. Chem. Rev. 104, 3607–3622 (2004).

3. Brutscher, B. et al. NMR methods for the study of instrinsically disordered proteins structure, dynamics, and interactions: General overview and practical guidelines. Intrinsically Disord. Proteins Studied by NMR Spectrosc. 49–122 (2015).

4. Jensen, M. R., Zweckstetter, M.Huang, J.-R. & Blackledge, M. Exploring free-energy landscapes of intrinsically disordered proteins at atomic resolution using NMR spectroscopy. Chem. Rev. 114, 6632–6660 (2014).

5. Yagasaki, M. & Hashimoto, S. Synthesis and application of dipeptides; current status and perspectives. Appl. Microbiol. Biotechnol. 81, 13–22 (2008).

6. Oh, K.-I. et al. A comprehensive library of blocked dipeptides reveals intrinsic backbone conformational propensities of unfolded proteins. Proteins: Struct. Funct. Bioinforma. 80, 977–990 (2012).

7. Uversky, V. N. & Dunker, A. K. Understanding protein non-folding. Biochimica et Biophys. Acta (BBA)-Proteins Proteomics 1804, 1231–1264 (2010).

8. Van Der Lee, R. et al. Classification of intrinsically disordered regions and proteins. Chem. Rev. 114, 6589–6631 (2014).

9. Dunker, A. K. et al. Intrinsically disordered protein. J. Mol. Graph. Model. 19, 26–59 (2001).

10. Babu, M. M. The contribution of intrinsically disordered regions to protein function, cellular complexity, and human disease. Biochem. Soc. Transactions 44, 1185–1200 (2016).

11. Oldfield, C. J. et al. Comparing and combining predictors of mostly disordered proteins. Biochemistry 44, 1989–2000 (2005).

12. Necci, M., Piovesan, D. & Tosatto, S. C. Critical assessment of protein intrinsic disorder prediction. Nat. Methods 18, 472–481 (2021).

13. Trivedi, R. & Nagarajaram, H. A. Intrinsically disordered proteins: An overview. Int. J. Mol. Sci. 23, 14050 (2022).

14. Evans, R., Ramisetty, S., Kulkarni, P. & Weninger, K. Illuminating intrinsically disordered proteins with integrative structural biology. Biomolecules 13, 124 (2023).

15. Maiti, S., Maji, T., Saibo, N. V., De, S. et al. Experimental methods to study the structure and dynamics of intrinsically disordered regions in proteins. Curr. Res. Struct. Biol. 100138 (2024).

16. Borthakur, K., Sisk, T. R., Panei, F. P., Bonomi, M. & Robustelli, P. Determining accurate conformational ensembles of intrinsically disordered proteins at atomic resolution. Nat. Commun. 16, 9036 (2025).

17. Kummerer, F., Orioli, S. & Lindorff-Larsen, K. Fitting force field parameters to NMR relaxation data. J. Chem. Theory Comput. 19, 3741–3751 (2023).

18. Rindfleisch, T. et al. Molecular dynamics of the intrinsically disordered protein COR15A—A force field validation on structure and dynamics. J. Chem. Theory Comput. 21, 9147–9163 (2025).

19. Ollila, O. S., Heikkinen, H. A. & Iwaï, H. Rotational dynamics of proteins from spin relaxation times and molecular dynamics simulations. The J. Phys. Chem. B 122, 6559–6569 (2018).

20. Salvi, N., Abyzov, A. & Blackledge, M. Multi-timescale dynamics in intrinsically disordered proteins from NMR relaxation and molecular simulation. The J. Phys. Chem. Lett. 7, 2483–2489 (2016).

21. Avbelj, F., Grdadolnik, S. G., Grdadolnik, J. & Baldwin, R. L. Intrinsic backbone preferences are fully present in blocked amino acids. Proc. Natl. Acad. Sci. 103, 1272–1277 (2006).

22. Eker, F., Cao, X., Nafie, L. & Schweitzer-Stenner, R. Tripeptides adopt stable structures in water. A combined polarized visible raman, FTIR, and VCD spectroscopy study. J. Am. Chem. Soc. 124, 14330–14341 (2002).

23. Shi, Z., Woody, R. W. & Kallenbach, N. R. Is polyproline II a major backbone conformation in unfolded proteins? Adv. Protein Chem. 62, 163–240 (2002).

24. Aspromonte, M. C. et al. DisProt in 2024: Improving function annotation of intrinsically disordered proteins. Nucleic Acids Res. 52, D434–D441 (2024).

25. Theillet, F.-X. et al. The alphabet of intrinsic disorder: I. Act like a Pro: On the abundance and roles of proline residues in intrinsically disordered proteins. Intrinsically Disord. Proteins 1, e24360 (2013).

26. Mateos, B. et al. The ambivalent role of proline residues in an intrinsically disordered protein: From disorder promoters to compaction facilitators. J. Mol. Biol. 432, 3093–3111 (2020).

27. Rindfleisch, T., Taule, E. F., Miettinen, M. S. & Underhaug, J. Data repository: Complete NMR assignment for 275 of the most common dipeptides in intrinsically disordered proteins. 10.17617/3.CBCNQG (2026). Edmond—The Open Research Data Repository of the Max Planck Society.

28. Hyberts, S. G., Milbradt, A. G., Wagner, A. B., Arthanari, H. & Wagner, G. Application of iterative soft thresholding for fast reconstruction of NMR data non-uniformly sampled with multidimensional Poisson Gap scheduling. J. Biomol. NMR 52, 315–327 (2012).

29. Pupier, M. et al. NMReDATA, a standard to report the NMR assignment and parameters of organic compounds. Magn. Reson. Chem. 56, 703–715 (2018).

30. Ulrich, E. L. et al. NMR-STAR: Comprehensive ontology for representing, archiving and exchanging data from nuclear magnetic resonance spectroscopic experiments. J. Biomol. NMR 73, 5–9 (2019).

31. Hoch, J. C. et al. Biological magnetic resonance data bank. Nucleic Acids Res. 51, D368–D376 (2023).

32. Mölder, F. et al. Sustainable data analysis with Snakemake. F1000Research 10, 33 (2025).

33. Dalby, A. et al. Description of several chemical structure file formats used by computer programs developed at molecular design limited. J. Chem. Inf. Comput. Sci. 32, 244–255 (1992).

34. Bernstein, F. C. et al. The protein data bank: A computer-based archival file for macromolecular structures. J. Mol. Biol. 112, 535–542 (1977).

35. Uversky, V. N. et al. Unfoldomics of human diseases: Linking protein intrinsic disorder with diseases. BMC Genomics 10, S7 (2009).

36. Ziarek, J. J., Baptista, D. & Wagner, G. Recent developments in solution nuclear magnetic resonance (NMR)-based molecular biology. J. Mol. Medicine 96, 1–8 (2018).

37. Wishart, D. S. Interpreting protein chemical shift data. Prog. Nucl. Magn. Reson. Spectrosc. 58, 62–87 (2011).

38. Granger, P., Bourdonneau, M., Assémat, O. & Piotto, M. NMR chemical shift measurements revisited: High precision measurements. Concepts Magn. Reson. Part A: An Educ. J. 30, 184–193 (2007).

39. Eggenberger, U., Pfäbdker, P. & Bodenhausen, G. Folding and pattern recognition in two-dimensional NMR spectra. J. Magn. Reson. 77, 192–196 (1988).

40. Wang, Y. & Jardetzky, O. Investigation of the neighboring residue effects on protein chemical shifts. J. Am. Chem. Soc. 124, 14075–14084 (2002).

41. Wang, B., Wang, Y. & Wishart, D. S. A probabilistic approach for validating protein NMR chemical shift assignments. J. Biomol. NMR 47, 85–99 (2010).

42. Klukowski, P., Riek, R. & Güntert, P. Rapid protein assignments and structures from raw NMR spectra with the deep learning technique ARTINA. Nat. Commun. 13, 6151 (2022).

43. Klukowski, P., Riek, R. & Güntert, P. Time-optimized protein NMR assignment with an integrative deep learning approach using AlphaFold and chemical shift prediction. Sci. Adv. 9, eadi9323 (2023).

44. Tamiola, K., Acar, B. & Mulder, F. A. Sequence-specific random coil chemical shifts of intrinsically disordered proteins. J. Am. Chem. Soc. 132, 18000–18003 (2010).

45. Nielsen, J. T. & Mulder, F. A. POTENCI: prediction of temperature, neighbor and pH-corrected chemical shifts for intrinsically disordered proteins. J. Biomol. NMR 70, 141–165 (2018).

46. Shen, Y., Delaglio, F., Cornilescu, G. & Bax, A. TALOS+: a hybrid method for predicting protein backbone torsion angles from NMR chemical shifts. J. Biomol. NMR 44, 213–223 (2009).

47. Shen, Y. & Bax, A. Protein structural information derived from NMR chemical shift with the neural network program TALOS-N. In Artificial Neural Networks, 17–32 (Springer, 2014).

48. Shen, Y. & Bax, A. SPARTA+: A modest improvement in empirical NMR chemical shift prediction by means of an artificial neural network. J. Biomol. NMR 48, 13–22 (2010).

49. Han, B., Liu, Y., Ginzinger, S. W. & Wishart, D. S. SHIFTX2: significantly improved protein chemical shift prediction. J. Biomol. NMR 50, 43–57 (2011).

50. Li, D.-W. & Brüschweiler, R. PPM: a side-chain and backbone chemical shift predictor for the assessment of protein conformational ensembles. J. Biomol. NMR 54, 257–265 (2012).

51. Lee, W., Tonelli, M. & Markley, J. L. NMRFAM-SPARKY: enhanced software for biomolecular NMR spectroscopy. Bioinformatics 31, 1325–1327 (2015).

52. Vranken, W. F. et al. The CCPN data model for NMR spectroscopy: Development of a software pipeline. Proteins: Struct. Funct. Bioinforma. 59, 687–696 (2005).

53. Fogh, R. et al. The CCPN project: an interim report on a data model for the nmr community. Nat. Struct. Biol. 9, 416–418 (2002).

54. Helmus, J. J. & Jaroniec, C. P. Nmrglue: An open source python package for the analysis of multidimensional NMR data. J. Biomol. NMR 55, 355–367 (2013).

