## Supplementary Information for "Complete NMR assignment for 275 of the most common dipeptides in intrinsically disordered proteins"

**Data availability**

The chemical shift datasets for 275 of the most common dipeptides in IDPs, along with the corresponding NMR spectra, are deposited in the Edmond Open Research Data Repository (edmond.mpg.de) at:

https://doi.org/10.17617/3.CBCNQG

### Supplementary Figures

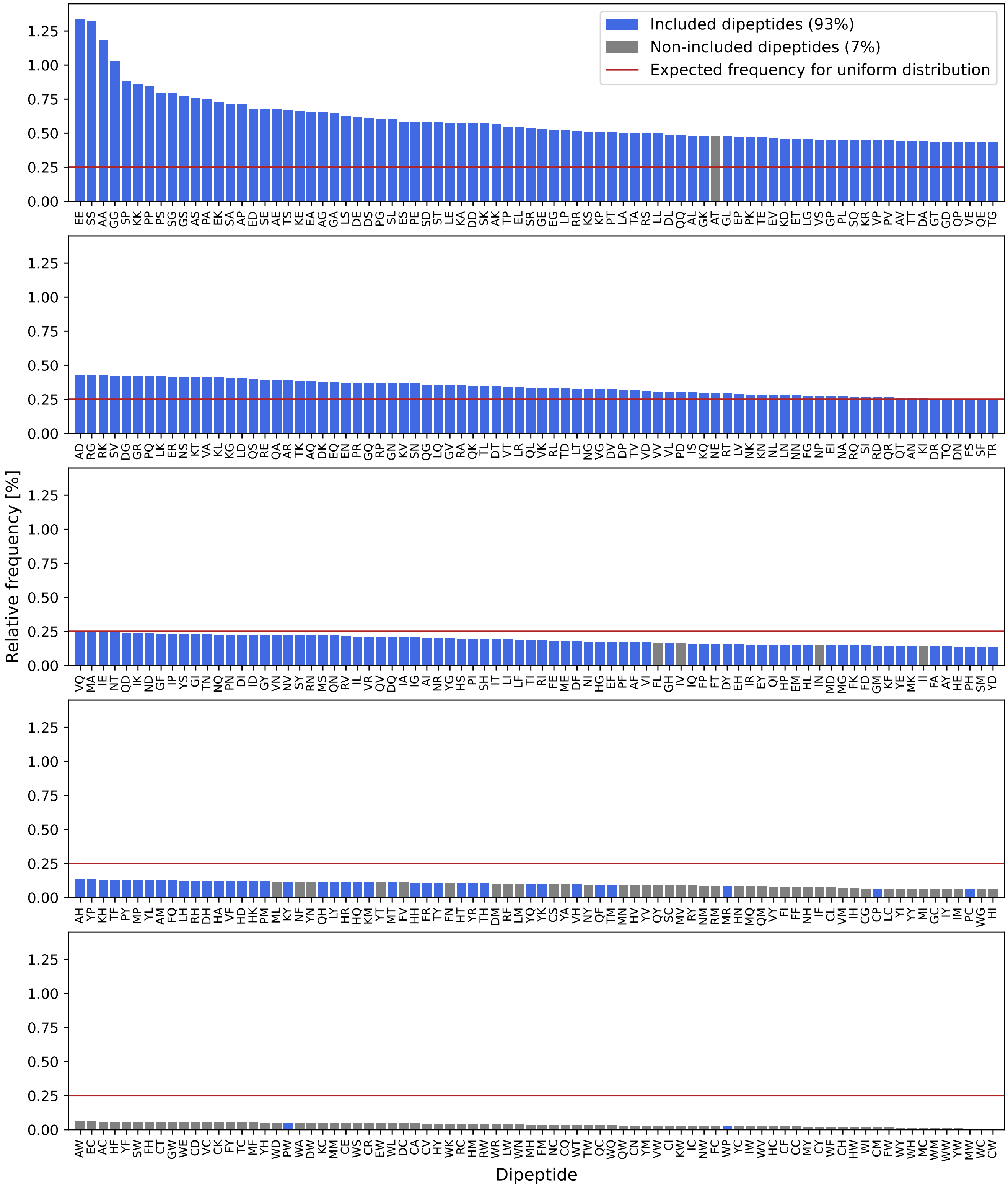


**Figure S1.** Relative frequencies of the 400 canonical dipeptides in the IDP proteome, as derived from the DisProt[1] database, compared to the uniform distribution (red line). Dipeptides included in the dataset are shown in blue and non-included ones in gray. The total frequency (%) is shown in the legend key for both populations. Additional details, including the relative and absolute frequencies, are provided in Table S2.

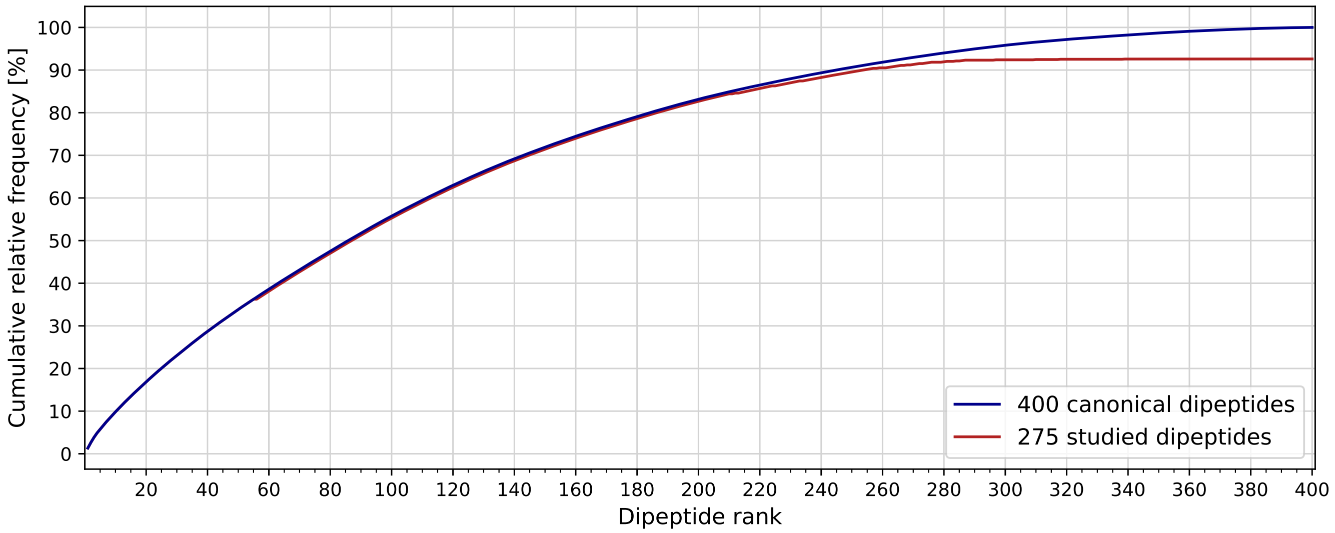


**Figure S2.** The cumulative distribution functions of the 275 studied (red), and all the 400 canonical dipeptides (blue) in the IDP proteome, derived from the DisProt[1] database, plotted as a function of the dipeptide rank (Figure 1 and Table S2).

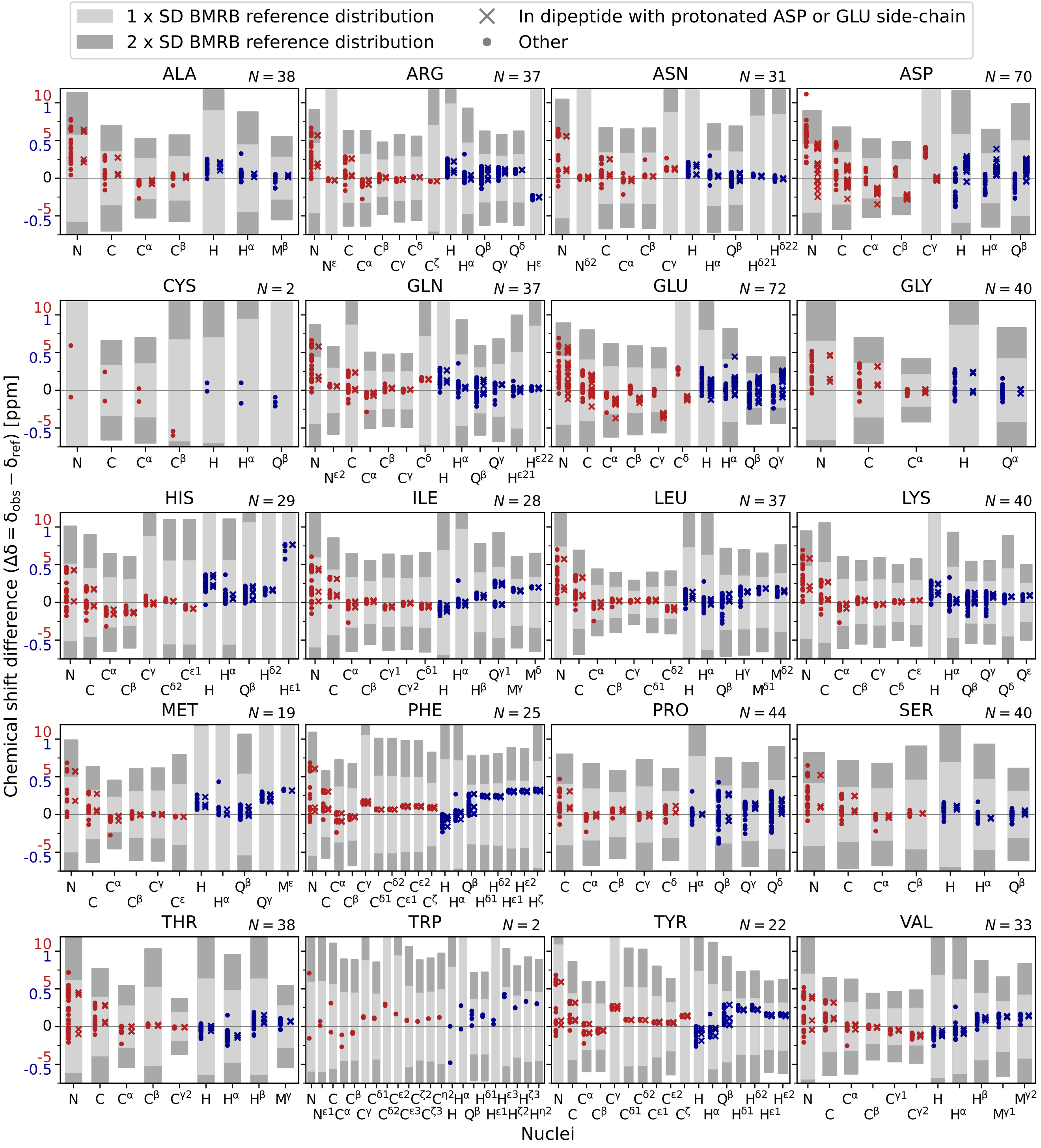


**Figure S3.** Residue-specific panels show the distributions of $\Delta\delta = \delta_{obs} - \delta_{ref}$ for each nucleus, defined as the difference between the observed chemical shifts ($\delta_{obs}$) and the BMRB reference values[2] ($\delta_{ref}$). Each panel has two $y$-axes, as the $^1$H $\Delta\delta$ (blue) span a different range than the $^{13}$C/$^{15}$N $\Delta\delta$ (red). Shaded regions indicate the BMRB reference standard deviation (SD) intervals[2]: ±1SD (light grey) and ±2SD (dark grey). $N$ is the total number of observations per residue; dipeptides with protonated and deprotonated ASP or GLU side-chains are considered separately. $^1$H resonances are classified as H, Q, and M: H represents CH protons with one data point; Q corresponds to $CH_2$ groups, for which the two protons are shown as separate data points; and M denotes methyl protons, generally yielding a single averaged chemical shift. Cross-symbols represent residues included in dipeptides with a protonated ASP or GLU side-chain, dots show residues in dipeptides with protonation states corresponding to those expected at pH 7 (ASP and GLU side-chains deprotonated).

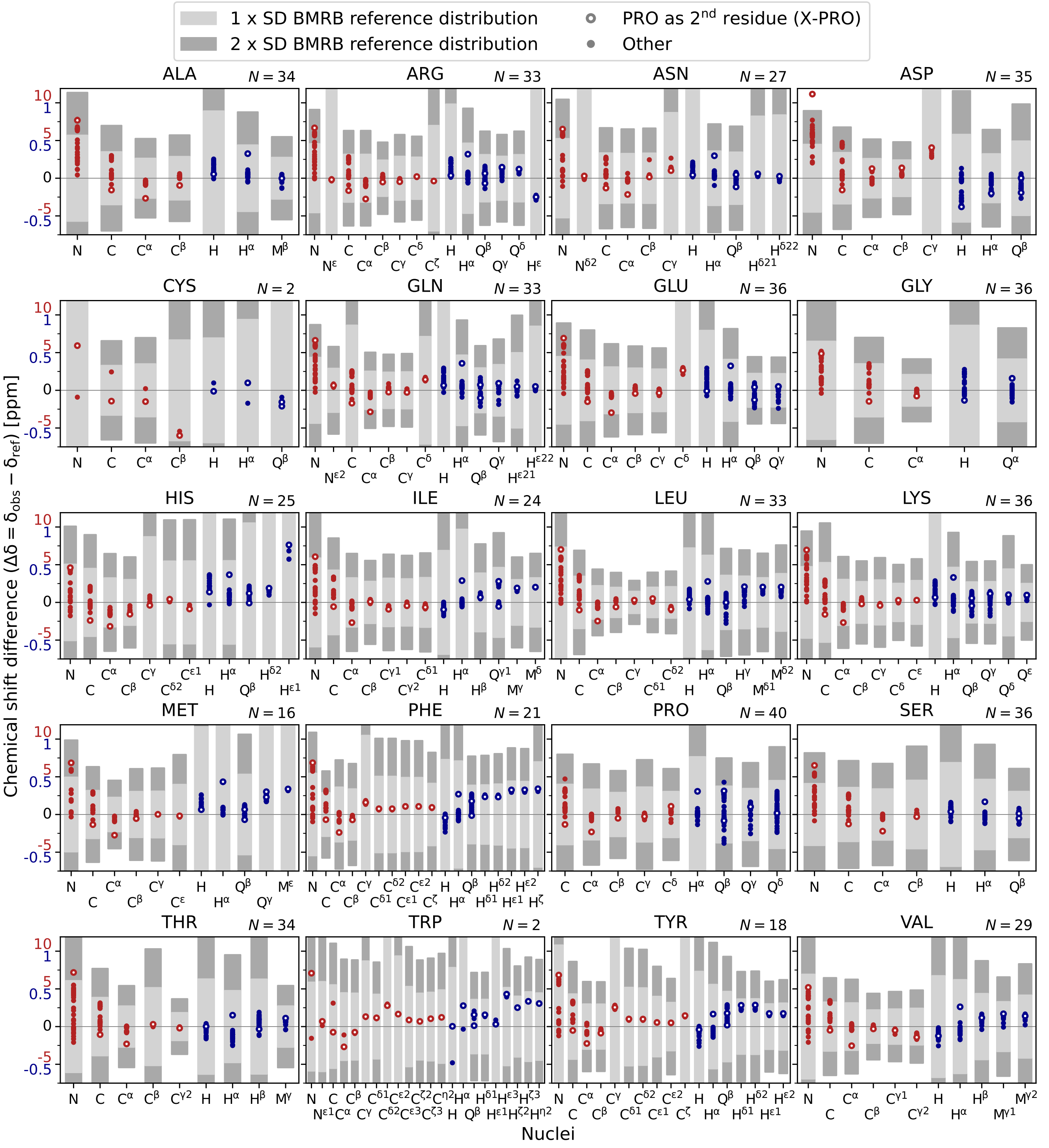


**Figure S4.** Dipeptides with protonation states corresponding to those expected at pH 7: Residue-specific panels show the distributions of $\Delta\delta = \delta_{obs} - \delta_{ref}$ for each nucleus, defined as the difference between the observed chemical shifts ($\delta_{obs}$) and the BMRB reference values[2] ($\delta_{ref}$). Each panel has two $y$-axes, as the $^{1}$H $\Delta\delta$ (blue) span a different range than the $^{13}$C/$^{15}$N $\Delta\delta$ (red). Shaded regions indicate the BMRB reference standard deviation (SD) intervals[2]: ±1SD (light grey) and ±2SD (dark grey). $N$ is the total number of observations per residue. $^{1}$H resonances are classified as H, Q, and M: H represents CH protons with one data point; Q corresponds to $CH_2$ groups, for which the two protons are shown as separate data points; and M denotes methyl protons, generally yielding a single averaged chemical shift. Circles indicate residues followed by proline (X–PRO motif) and dots represent all others.

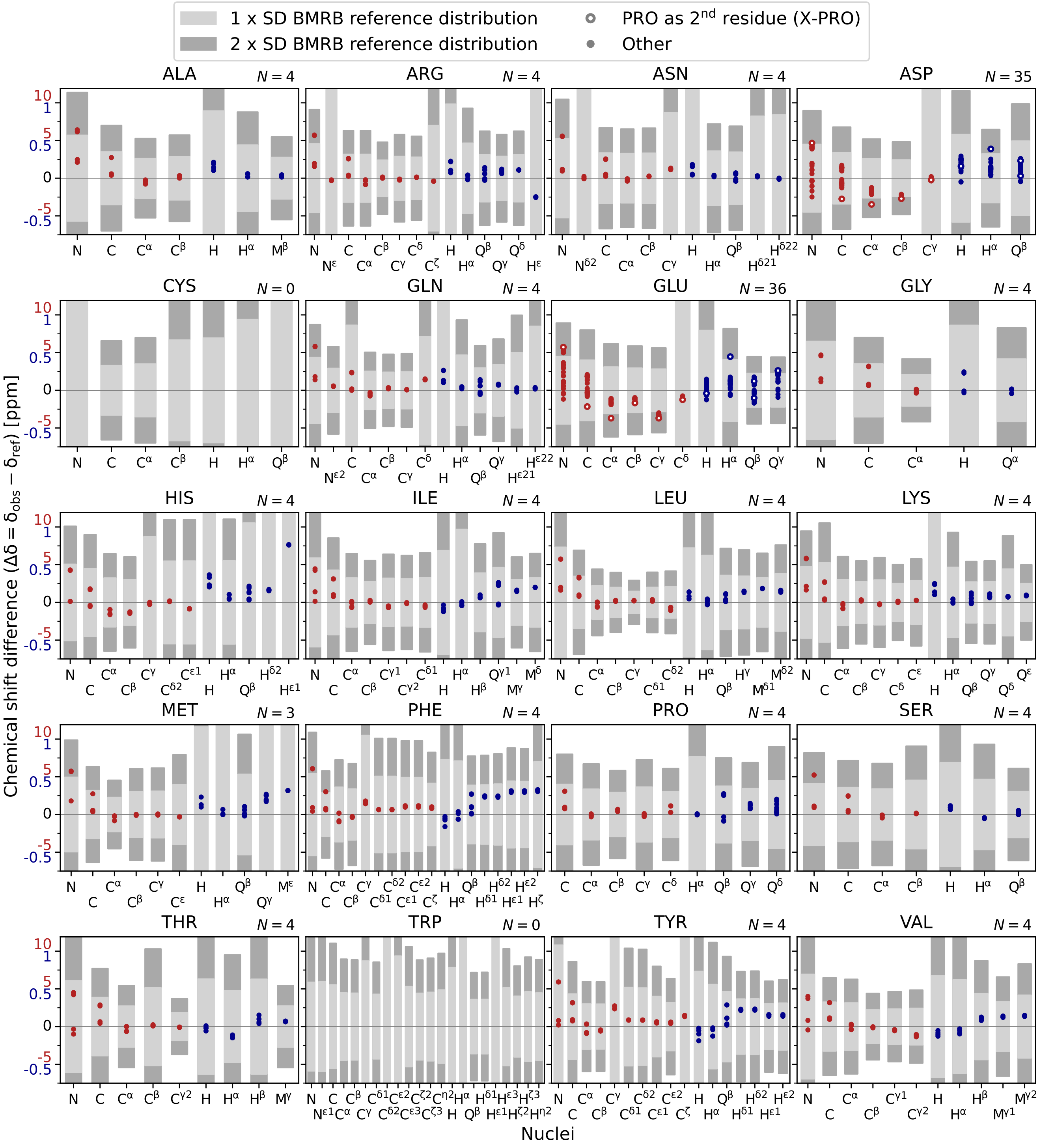


**Figure S5.** ASP or GLU containing dipeptides with protonated side-chains: Residue-specific panels show the distributions of $\Delta\delta = \delta_{obs} - \delta_{ref}$ for each nucleus, defined as the difference between the observed chemical shifts ($\delta_{obs}$) and the BMRB reference values[2] ($\delta_{ref}$). Each panel has two *y*-axes, as the $^{1}$H $\Delta\delta$ (blue) span a different range than the $^{13}$C/$^{15}$N $\Delta\delta$ (red). Shaded regions indicate the BMRB reference standard deviation (SD) intervals[2]: $\pm$1SD (light grey) and $\pm$2SD (dark grey). *N* is the total number of observations per residue. $^{1}$H resonances are classified as H, Q, and M: H represents CH protons with one data point; Q corresponds to $CH_2$ groups, for which the two protons are shown as separate data points; and M denotes methyl protons, generally yielding a single averaged chemical shift. Circles indicate residues followed by proline (X–PRO motif) and dots represent all others.

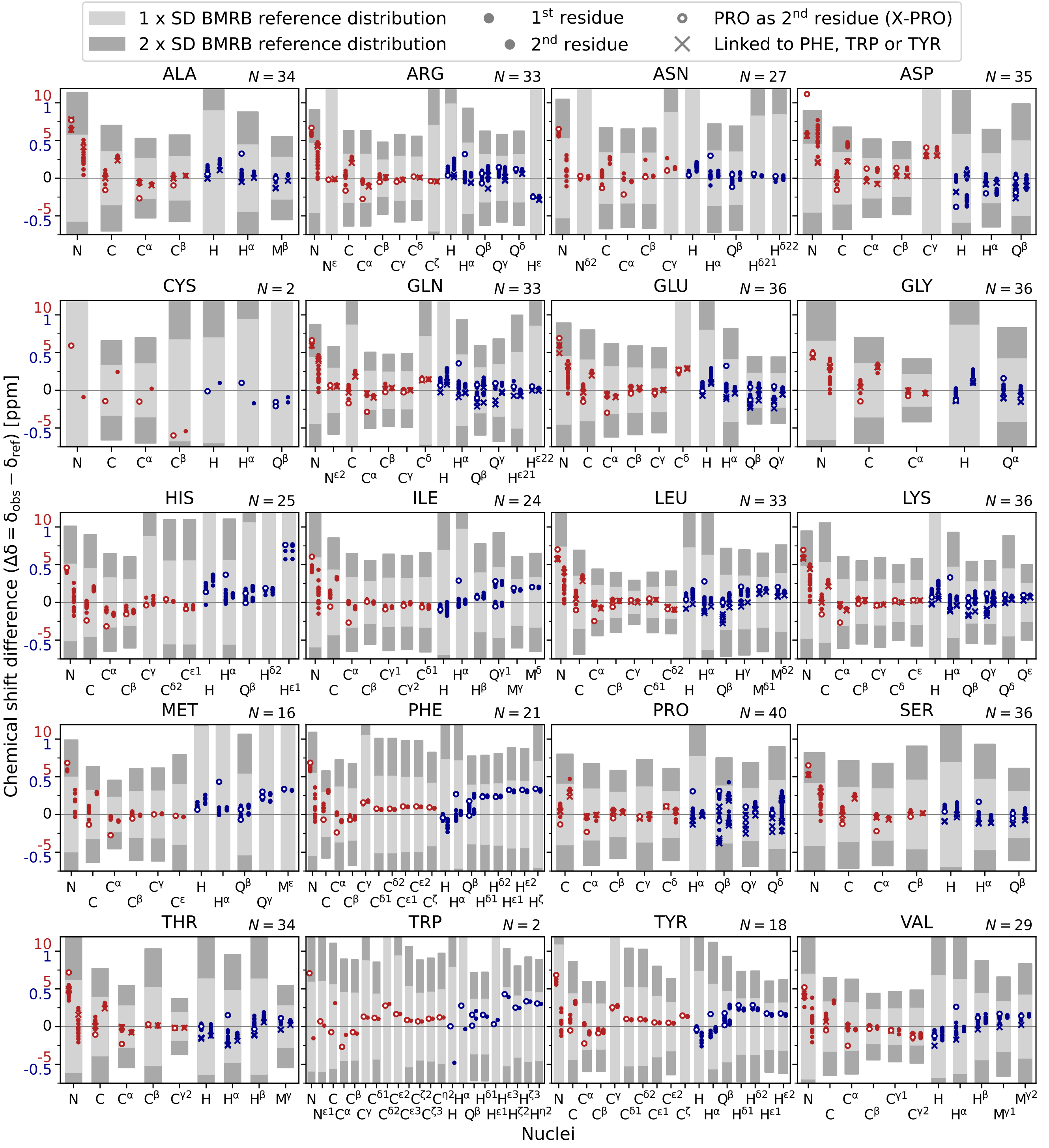


**Figure S6.** Dipeptides with protonation states corresponding to those expected at pH 7: Residue-specific panels display the distributions of $\Delta\delta = \delta_{obs} - \delta_{ref}$ for each nucleus, defined as the difference between the observed chemical shifts ($\delta_{obs}$) and the BMRB reference values[2] ($\delta_{ref}$). Each panel includes two $y$-axes, as the $^{1}$H $\Delta\delta$ (blue) span a narrower range than the $^{13}$C/$^{15}$N $\Delta\delta$ (red). Shaded regions mark the BMRB reference standard deviation (SD) intervals[2]: ±1SD (light grey) and ±2SD (dark grey). $N$ is the total number of observations per residue. $^{1}$H resonances are classified as H, Q, and M: H represents CH protons with one data point; Q corresponds to $CH_2$ groups, for which the two protons are shown as separate data points; and M denotes methyl protons, generally yielding a single averaged chemical shift. Data points corresponding to residues in the first position of the dipeptide are plotted on the left, while those in the second position are plotted on the right. Circles indicate residues followed by proline (X–PRO motif), dots represent all others, and cross-symbols mark residues linked to aromatic residues (PHE, TRP, or TYR).

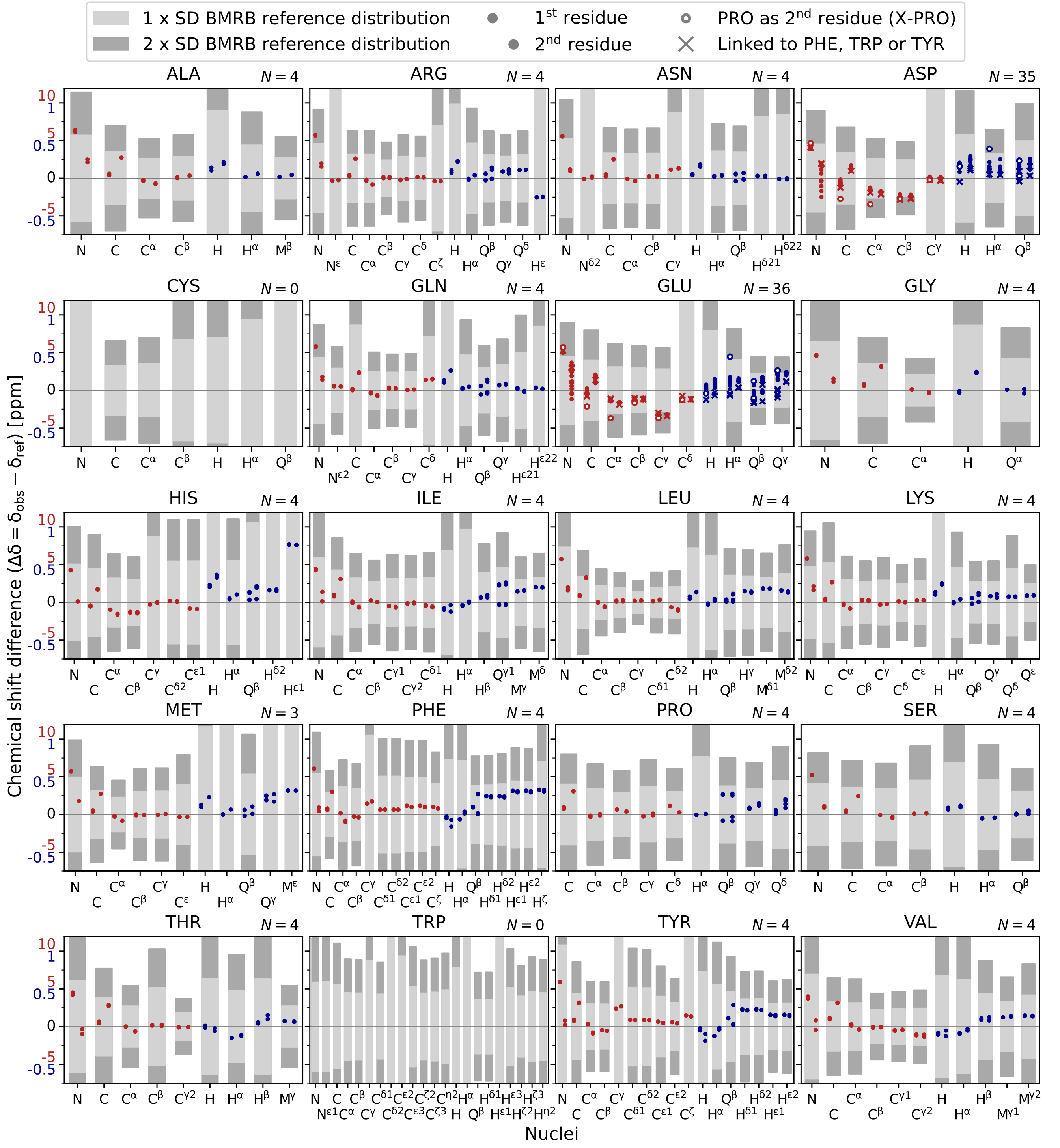


**Figure S7.** ASP or GLU containing dipeptides with protonated side-chains: Residue-specific panels display the distributions of $\Delta\delta = \delta_{obs} - \delta_{ref}$ for each nucleus, defined as the difference between the observed chemical shifts ($\delta_{obs}$) and the BMRB reference values[2] ($\delta_{ref}$). Each panel includes two $y$-axes, as the $^{1}$H $\Delta\delta$ (blue) span a narrower range than the $^{13}$C/$^{15}$N $\Delta\delta$ (red). Shaded regions mark the BMRB reference standard deviation (SD) intervals[2]: $\pm$1SD (light grey) and $\pm$2SD (dark grey). $N$ is the total number of observations per residue. $^{1}$H resonances are classified as H, Q, and M: H represents CH protons with one data point; Q corresponds to $CH_2$ groups, for which the two protons are shown as separate data points; and M denotes methyl protons, generally yielding a single averaged chemical shift. Data points corresponding to residues in the first position of the dipeptide are plotted on the left, while those in the second position are plotted on the right. Circles indicate residues followed by proline (X–PRO motif), dots represent all others, and cross-symbols mark residues linked to aromatic residues (PHE, TRP, or TYR).

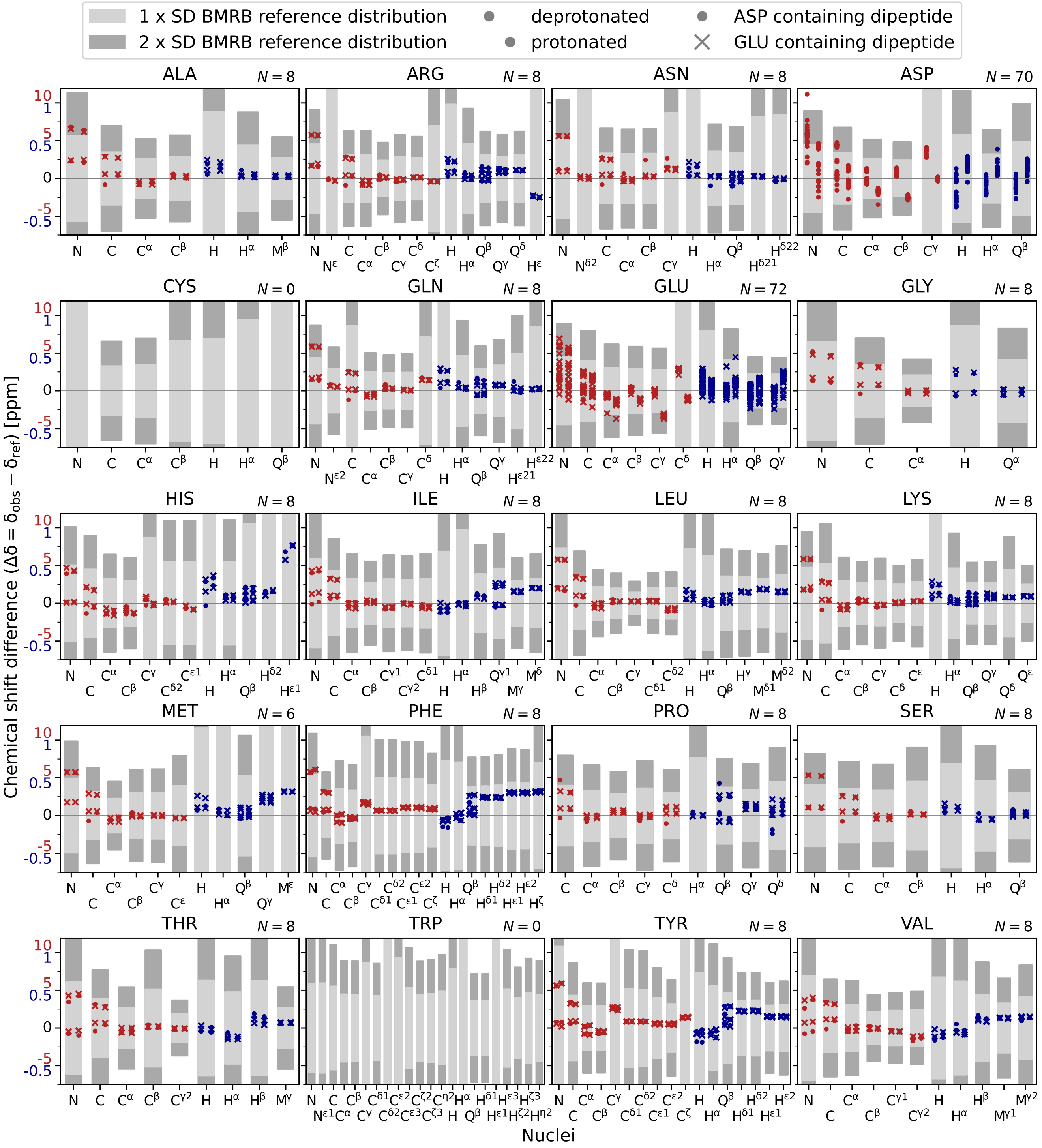


**Figure S8.** Protonation effects in ASP and GLU containing dipeptides: Residue-specific panels display the distributions of $\Delta\delta = \delta_{obs} - \delta_{ref}$ for each nucleus, defined as the difference between the observed chemical shifts ($\delta_{obs}$) and the BMRB reference values[2] ($\delta_{ref}$). Each panel includes two $y$-axes, as the $^{1}$H $\Delta\delta$ (blue) span a narrower range than the $^{13}$C/$^{15}$N $\Delta\delta$ (red). Shaded regions mark the BMRB reference standard deviation (SD) intervals[2]: ±1SD (light grey) and ±2SD (dark grey). $N$ is the total number of observations per residue. $^{1}$H resonances are classified as H, Q, and M: H represents CH protons with one data point; Q corresponds to $CH_2$ groups, for which the two protons are shown as separate data points; and M denotes methyl protons, generally yielding a single averaged chemical shift. Data points corresponding to dipeptides with deprotonated GLU/ASP are plotted on the left, while those with protonated GLU/ASP are plotted on the right. Circles indicate ASP-containing dipeptides and cross-symbols mark dipeptides including GLU.

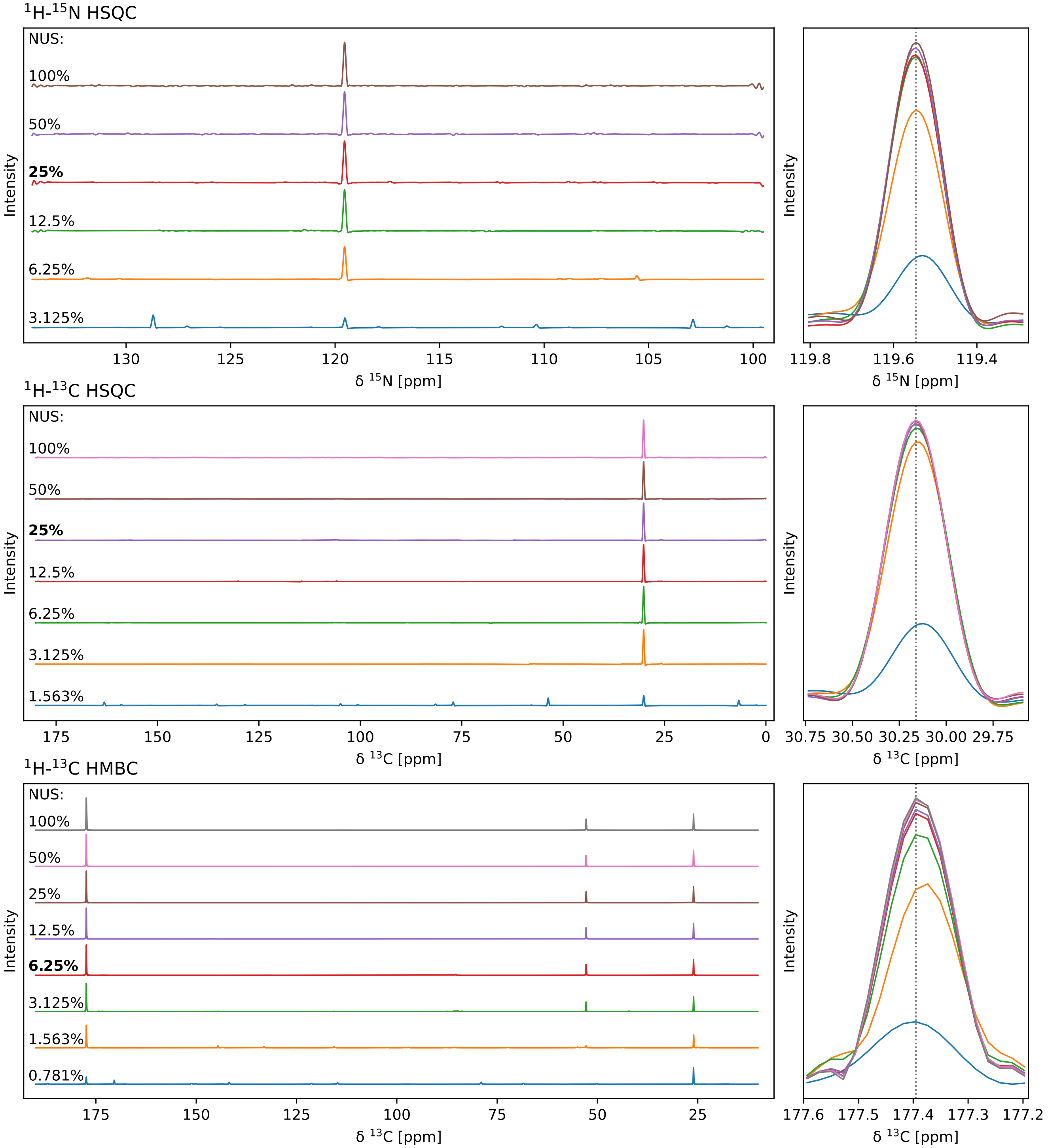


**Figure S9.** Assessment of NUS-related effects, exemplarily demonstrated for the dipeptide AE. To evaluate potential artifact occurrence (left panels) and perturbations of chemical shift ($\delta$) positions (right panels), we show 1D columns extracted from the corresponding 2D spectra at $^{1}$H = 8.38 ppm for the $^{1}$H–$^{15}$N HSQC, and at $^{1}$H = 2.48 ppm for the $^{1}$H–$^{13}$C HSQC and $^{1}$H–$^{13}$C HMBC, using NUS fractions from 0.781% to 100%. The results show no effect on chemical-shift positions for the NUS fractions used in this study (highlighted in bold). Furthermore, peak intensities at 100% sampling (uniform sampling) and at the applied NUS fractions differ only marginally, and no spectral artifacts are observed, except for a single marginal artificial peak in the HMBC dataset.

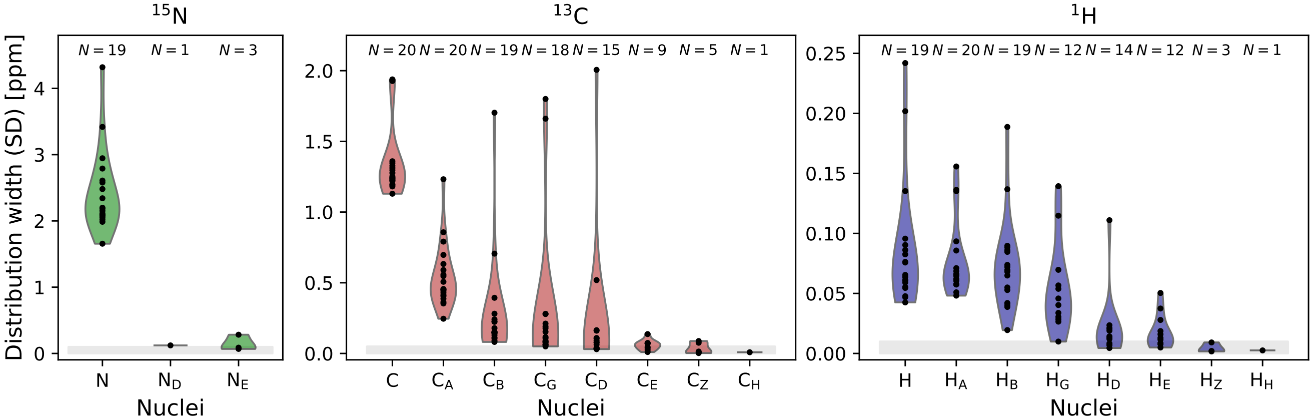


**Figure S10.** For every residue–atom-type combination plotted in Figure 2 (main text), the standard deviation ($SD_{res,at}$, with res: residue and at: atom type) of the corresponding data points (Figure 2) was calculated and is shown as a black dot. The resulting $SD_{res,at}$ values were then grouped according to the atom type ($x$-axis) and are plotted separately for $^{15}$N, $^{13}$C, and $^{1}$H. For every atom type, the underlying distributions of computed $SD_{res,at}$-values are visualized as violins, with $N$ indicating the number of individual observations per distribution. Broader distributions or higher $SD_{res,at}$ values reflect greater chemical shift variability for the corresponding atom types. Gray-shaded regions indicate the typical assignment uncertainties of the respective nuclei ($^{15}$N: 0.1 ppm, $^{13}$C: 0.05 ppm, and $^{1}$H: 0.01 ppm).

### Supplementary Tables

**Table S1.** Overview of NMR instruments used for individual dipeptide measurements.

| Instrument | Analyzed dipeptides |
|---|---|
| Ascend 600 MHz with QCI-P CryoProbe | AA, AD, AE, AG, AK, AL, AN, AP, AQ, AR, AS, AV, CP, DA, DD, DE, DG, DH, DI, DK, DL, DN, DP, DQ, DR, DS, DT, DV, EA, ED, EE, EF, EG, EH, EI, EK, EL, EN, EP, EQ, ER, ES, ET, EV, EY, FA, FG, FK, FP, FS, GA, GD, GE, GF, GG, GH, GI, GK, GL, GN, GP, GQ, GR, GS, GT, GV, GY, HE, HG, HH, HK, HL, HP, HQ, IA, ID, IE, IG, IK, IL, IP, IS, KA, KD, KE, KF, KG, KH, KI, KK, KL, KN, KP, KQ, KR, KS, KT, KV, LA, LD, LE, LF, LG, LI, LK, LL, LN, LP, LQ, LR, LS, LT, LV, MA, MD, ME, MG, MP, NA, ND, NE, NG, NK, NL, NP, NS, NT, NV, PA, PC, PD, PE, PF, PG, PH, PI, PK, PL, PN, PP, PQ, PR, PS, PT, PW, PY, QA, QD, QE, QG, QK, QL, QN, QP, QR, QS, QT, QV, RA, RD, RE, RG, RK, RL, RP, RQ, RS, RT, SA, SD, SE, SF, SG, SH, SI, SK, SL, SN, SP, SQ, SR, SS, ST, SV, SY, TA, TD, TE, TG, TH, TK, TL, TM, TN, TP, TQ, TR, TS, TT, TV, VA, VD, VE, VF, VG, VH, VK, VL, VP, VQ, VR, VT, VV, YE, YG, YP |
| Ascend 850 MHz with TCI CryoProbe | AF, AH, AI, AM, AY, DF, DY, EM, FD, FE, FQ, FR, FT, GM, HA, HD, HR, HS, HT, IQ, IR, IT, KM, KY, LH, LY, MK, MR, MS, MT, NN, NQ, NR, PM, PV, QF, QH, QI, RH, RI, RN, RR, RV, SM, TF, TI, TY, VI, VN, WP, YD, YK, YL, YQ, YR, YS |

**Table S2.** The relative frequencies (rel. freq.) of all the 400 canonical dipeptides in the DisProt[1] database (that contains experimentally-confirmed IDP sequences); their absolute occurrence (abs. occurr.) in DisProt; and their relative frequency in the dataset presented here (dipeptides not included are indicated as "–"). Entries are sorted by the relative frequency / rank (left; according to Figure 1) and by the dipeptide in alphabetical order (right).

| Sorted by relative frequency (rank) | | | | | Sorted alphabetically by dipeptide name | | | | |
|---|---|---|---|---|---|---|---|---|---|
| Rank | Dipeptide | Rel. freq. in DisProt [%] | Abs. occurr. in DisProt | Rel. freq. in dataset [%] | Rank | Dipeptide | Rel. freq. in DisProt [%] | Abs. occurr. in DisProt | Rel. freq. in dataset [%] |
| 1 | EE | 1.3333 | 2955 | 1.4397 | 3 | AA | 1.1866 | 2630 | 1.2813 |
| 2 | SS | 1.3224 | 2931 | 1.4280 | 323 | AC | 0.0559 | 124 | – |
| 3 | AA | 1.1866 | 2630 | 1.2813 | 81 | AD | 0.4309 | 955 | 0.4653 |
| 4 | GG | 1.0296 | 2282 | 1.1118 | 18 | AE | 0.6768 | 1500 | 0.7308 |
| 5 | SP | 0.8816 | 1954 | 0.9520 | 209 | AF | 0.1696 | 376 | 0.1832 |
| 6 | KK | 0.8636 | 1914 | 0.9325 | 22 | AG | 0.6520 | 1445 | 0.7040 |
| 7 | PP | 0.8464 | 1876 | 0.9140 | 241 | AH | 0.1331 | 295 | 0.1437 |
| 8 | PS | 0.7995 | 1772 | 0.8633 | 191 | AI | 0.2008 | 445 | 0.2168 |
| 9 | SG | 0.7927 | 1757 | 0.8560 | 37 | AK | 0.5653 | 1253 | 0.6105 |
| 10 | GS | 0.7706 | 1708 | 0.8321 | 54 | AL | 0.4778 | 1059 | 0.5159 |
| 11 | AS | 0.7553 | 1674 | 0.8156 | 248 | AM | 0.1263 | 280 | 0.1364 |
| 12 | PA | 0.7508 | 1664 | 0.8107 | 153 | AN | 0.2585 | 573 | 0.2792 |
| 13 | EK | 0.7269 | 1611 | 0.7849 | 15 | AP | 0.7129 | 1580 | 0.7698 |
| 14 | SA | 0.7178 | 1591 | 0.7751 | 101 | AQ | 0.3853 | 854 | 0.4161 |
| 15 | AP | 0.7129 | 1580 | 0.7698 | 99 | AR | 0.3907 | 866 | 0.4219 |
| 16 | ED | 0.6808 | 1509 | 0.7352 | 11 | AS | 0.7553 | 1674 | 0.8156 |
| 17 | SE | 0.6786 | 1504 | 0.7328 | 56 | AT | 0.4765 | 1056 | – |
| 18 | AE | 0.6768 | 1500 | 0.7308 | 72 | AV | 0.4422 | 980 | 0.4775 |
| 19 | TS | 0.6687 | 1482 | 0.7220 | 321 | AW | 0.0591 | 131 | – |
| 20 | KE | 0.6632 | 1470 | 0.7162 | 236 | AY | 0.1376 | 305 | 0.1486 |
| 21 | EA | 0.6578 | 1458 | 0.7103 | 350 | CA | 0.0456 | 101 | – |
| 22 | AG | 0.6520 | 1445 | 0.7040 | 383 | CC | 0.0230 | 51 | – |
| 23 | GA | 0.6461 | 1432 | 0.6977 | 331 | CD | 0.0514 | 114 | – |
| 24 | LS | 0.6235 | 1382 | 0.6733 | 344 | CE | 0.0474 | 105 | – |

Continued on next page

**Table S2 (continued)**

| Sorted by relative frequency (rank) | | | | | Sorted alphabetically by dipeptide name | | | | |
|---|---|---|---|---|---|---|---|---|---|
| Rank | Dipep-tide | Rel. freq. in DisProt [%] | Abs. occurr. in DisProt | Rel. freq. in dataset [%] | Rank | Dipep-tide | Rel. freq. in DisProt [%] | Abs. occurr. in DisProt | Rel. freq. in dataset [%] |
| 25 | DE | 0.6208 | 1376 | 0.6704 | 382 | CF | 0.0235 | 52 | – |
| 26 | DS | 0.6091 | 1350 | 0.6577 | 309 | CG | 0.0663 | 147 | – |
| 27 | PG | 0.6087 | 1349 | 0.6572 | 387 | CH | 0.0180 | 40 | – |
| 28 | SL | 0.6055 | 1342 | 0.6538 | 372 | CI | 0.0298 | 66 | – |
| 29 | ES | 0.5865 | 1300 | 0.6334 | 333 | CK | 0.0510 | 113 | – |
| 30 | PE | 0.5865 | 1300 | 0.6334 | 306 | CL | 0.0735 | 163 | – |
| 31 | SD | 0.5843 | 1295 | 0.6309 | 390 | CM | 0.0162 | 36 | – |
| 32 | ST | 0.5811 | 1288 | 0.6275 | 369 | CN | 0.0307 | 68 | – |
| 33 | LE | 0.5753 | 1275 | 0.6212 | 310 | CP | 0.0659 | 146 | 0.0711 |
| 34 | KA | 0.5730 | 1270 | 0.6187 | 363 | CQ | 0.0334 | 74 | – |
| 35 | DD | 0.5712 | 1266 | 0.6168 | 346 | CR | 0.0465 | 103 | – |
| 36 | SK | 0.5708 | 1265 | 0.6163 | 282 | CS | 0.0993 | 220 | – |
| 37 | AK | 0.5653 | 1253 | 0.6105 | 328 | CT | 0.0523 | 116 | – |
| 38 | TP | 0.5473 | 1213 | 0.5910 | 351 | CV | 0.0442 | 98 | – |
| 39 | EL | 0.5455 | 1209 | 0.5890 | 400 | CW | 0.0054 | 12 | – |
| 40 | SR | 0.5369 | 1190 | 0.5798 | 385 | CY | 0.0217 | 48 | – |
| 41 | GE | 0.5301 | 1175 | 0.5725 | 74 | DA | 0.4404 | 976 | 0.4755 |
| 42 | EG | 0.5247 | 1163 | 0.5666 | 349 | DC | 0.0456 | 101 | – |
| 43 | LP | 0.5211 | 1155 | 0.5627 | 35 | DD | 0.5712 | 1266 | 0.6168 |
| 44 | RR | 0.5175 | 1147 | 0.5588 | 25 | DE | 0.6208 | 1376 | 0.6704 |
| 45 | KS | 0.5098 | 1130 | 0.5505 | 204 | DF | 0.1778 | 394 | 0.1920 |
| 46 | KP | 0.5089 | 1128 | 0.5496 | 85 | DG | 0.4214 | 934 | 0.4550 |
| 47 | PT | 0.5076 | 1125 | 0.5481 | 252 | DH | 0.1223 | 271 | 0.1320 |
| 48 | LA | 0.5035 | 1116 | 0.5437 | 175 | DI | 0.2229 | 494 | 0.2407 |
| 49 | TA | 0.5004 | 1109 | 0.5403 | 102 | DK | 0.3804 | 843 | 0.4107 |
| 50 | RS | 0.4986 | 1105 | 0.5384 | 52 | DL | 0.4859 | 1077 | 0.5247 |
| 51 | LL | 0.4986 | 1105 | 0.5384 | 277 | DM | 0.1024 | 227 | – |
| 52 | DL | 0.4859 | 1077 | 0.5247 | 157 | DN | 0.2527 | 560 | 0.2728 |
| 53 | QQ | 0.4846 | 1074 | 0.5233 | 128 | DP | 0.3203 | 710 | 0.3459 |
| 54 | AL | 0.4778 | 1059 | 0.5159 | 188 | DQ | 0.2071 | 459 | 0.2236 |
| 55 | GK | 0.4774 | 1058 | 0.5155 | 155 | DR | 0.2545 | 564 | 0.2748 |
| 56 | AT | 0.4765 | 1056 | – | 26 | DS | 0.6091 | 1350 | 0.6577 |
| 57 | GL | 0.4746 | 1052 | 0.5125 | 117 | DT | 0.3479 | 771 | 0.3756 |
| 58 | EP | 0.4742 | 1051 | 0.5121 | 127 | DV | 0.3244 | 719 | 0.3503 |
| 59 | PK | 0.4724 | 1047 | 0.5101 | 341 | DW | 0.0487 | 108 | – |
| 60 | TE | 0.4715 | 1045 | 0.5091 | 217 | DY | 0.1552 | 344 | 0.1676 |
| 61 | EV | 0.4620 | 1024 | 0.4989 | 21 | EA | 0.6578 | 1458 | 0.7103 |
| 62 | KD | 0.4598 | 1019 | 0.4965 | 322 | EC | 0.0591 | 131 | – |
| 63 | ET | 0.4580 | 1015 | 0.4945 | 16 | ED | 0.6808 | 1509 | 0.7352 |
| 64 | LG | 0.4575 | 1014 | 0.4940 | 1 | EE | 1.3333 | 2955 | 1.4397 |
| 65 | VS | 0.4521 | 1002 | 0.4882 | 207 | EF | 0.1701 | 377 | 0.1837 |
| 66 | GP | 0.4507 | 999 | 0.4867 | 42 | EG | 0.5247 | 1163 | 0.5666 |
| 67 | PL | 0.4503 | 998 | 0.4862 | 218 | EH | 0.1552 | 344 | 0.1676 |
| 68 | SQ | 0.4489 | 995 | 0.4848 | 146 | EI | 0.2721 | 603 | 0.2938 |
| 69 | KR | 0.4471 | 991 | 0.4828 | 13 | EK | 0.7269 | 1611 | 0.7849 |
| 70 | VP | 0.4467 | 990 | 0.4823 | 39 | EL | 0.5455 | 1209 | 0.5890 |
| 71 | PV | 0.4467 | 990 | 0.4823 | 223 | EM | 0.1502 | 333 | 0.1622 |
| 72 | AV | 0.4422 | 980 | 0.4775 | 104 | EN | 0.3722 | 825 | 0.4019 |
| 73 | TT | 0.4408 | 977 | 0.4760 | 58 | EP | 0.4742 | 1051 | 0.5121 |

Continued on next page

**Table S2 (continued)**

| Sorted by relative frequency (rank) | | | | | Sorted alphabetically by dipeptide name | | | | |
|---|---|---|---|---|---|---|---|---|---|
| Rank | Dipep-tide | Rel. freq. in DisProt [%] | Abs. occurr. in DisProt | Rel. freq. in dataset [%] | Rank | Dipep-tide | Rel. freq. in DisProt [%] | Abs. occurr. in DisProt | Rel. freq. in dataset [%] |
| 74 | DA | 0.4404 | 976 | 0.4755 | 103 | EQ | 0.3781 | 838 | 0.4083 |
| 75 | GT | 0.4345 | 963 | 0.4692 | 89 | ER | 0.4160 | 922 | 0.4492 |
| 76 | GD | 0.4345 | 963 | 0.4692 | 29 | ES | 0.5865 | 1300 | 0.6334 |
| 77 | QP | 0.4345 | 963 | 0.4692 | 63 | ET | 0.4580 | 1015 | 0.4945 |
| 78 | VE | 0.4340 | 962 | 0.4687 | 61 | EV | 0.4620 | 1024 | 0.4989 |
| 79 | QE | 0.4340 | 962 | 0.4687 | 347 | EW | 0.0465 | 103 | – |
| 80 | TG | 0.4327 | 959 | 0.4672 | 220 | EY | 0.1539 | 341 | 0.1661 |
| 81 | AD | 0.4309 | 955 | 0.4653 | 235 | FA | 0.1394 | 309 | 0.1505 |
| 82 | RG | 0.4277 | 948 | 0.4619 | 376 | FC | 0.0266 | 59 | – |
| 83 | RK | 0.4264 | 945 | 0.4604 | 229 | FD | 0.1462 | 324 | 0.1579 |
| 84 | SV | 0.4232 | 938 | 0.4570 | 202 | FE | 0.1814 | 402 | 0.1959 |
| 85 | DG | 0.4214 | 934 | 0.4550 | 303 | FF | 0.0790 | 175 | – |
| 86 | GR | 0.4210 | 933 | 0.4546 | 144 | FG | 0.2743 | 608 | 0.2962 |
| 87 | PQ | 0.4210 | 933 | 0.4546 | 327 | FH | 0.0528 | 117 | – |
| 88 | LK | 0.4196 | 930 | 0.4531 | 302 | FI | 0.0794 | 176 | – |
| 89 | ER | 0.4160 | 922 | 0.4492 | 228 | FK | 0.1462 | 324 | 0.1579 |
| 90 | NS | 0.4133 | 916 | 0.4463 | 211 | FL | 0.1678 | 372 | – |
| 91 | KT | 0.4115 | 912 | 0.4443 | 361 | FM | 0.0356 | 79 | – |
| 92 | VA | 0.4115 | 912 | 0.4443 | 273 | FN | 0.1056 | 234 | – |
| 93 | KL | 0.4115 | 912 | 0.4443 | 215 | FP | 0.1588 | 352 | 0.1715 |
| 94 | KG | 0.4088 | 906 | 0.4414 | 249 | FQ | 0.1236 | 274 | 0.1335 |
| 95 | LD | 0.4083 | 905 | 0.4409 | 271 | FR | 0.1087 | 241 | 0.1174 |
| 96 | QS | 0.3961 | 878 | 0.4278 | 158 | FS | 0.2495 | 553 | 0.2694 |
| 97 | RE | 0.3948 | 875 | 0.4263 | 216 | FT | 0.1557 | 345 | 0.1681 |
| 98 | QA | 0.3912 | 867 | 0.4224 | 269 | FV | 0.1101 | 244 | – |
| 99 | AR | 0.3907 | 866 | 0.4219 | 391 | FW | 0.0149 | 33 | – |
| 100 | TK | 0.3853 | 854 | 0.4161 | 334 | FY | 0.0510 | 113 | – |
| 101 | AQ | 0.3853 | 854 | 0.4161 | 23 | GA | 0.6461 | 1432 | 0.6977 |
| 102 | DK | 0.3804 | 843 | 0.4107 | 315 | GC | 0.0627 | 139 | – |
| 103 | EQ | 0.3781 | 838 | 0.4083 | 76 | GD | 0.4345 | 963 | 0.4692 |
| 104 | EN | 0.3722 | 825 | 0.4019 | 41 | GE | 0.5301 | 1175 | 0.5725 |
| 105 | PR | 0.3713 | 823 | 0.4010 | 168 | GF | 0.2328 | 516 | 0.2514 |
| 106 | GQ | 0.3691 | 818 | 0.3985 | 4 | GG | 1.0296 | 2282 | 1.1118 |
| 107 | RP | 0.3673 | 814 | 0.3966 | 212 | GH | 0.1674 | 371 | 0.1808 |
| 108 | GN | 0.3673 | 814 | 0.3966 | 171 | GI | 0.2306 | 511 | 0.2490 |
| 109 | KV | 0.3659 | 811 | 0.3951 | 55 | GK | 0.4774 | 1058 | 0.5155 |
| 110 | SN | 0.3659 | 811 | 0.3951 | 57 | GL | 0.4746 | 1052 | 0.5125 |
| 111 | QG | 0.3582 | 794 | 0.3868 | 230 | GM | 0.1457 | 323 | 0.1574 |
| 112 | LQ | 0.3578 | 793 | 0.3864 | 108 | GN | 0.3673 | 814 | 0.3966 |
| 113 | GV | 0.3564 | 790 | 0.3849 | 66 | GP | 0.4507 | 999 | 0.4867 |
| 114 | RA | 0.3546 | 786 | 0.3829 | 106 | GQ | 0.3691 | 818 | 0.3985 |
| 115 | QK | 0.3492 | 774 | 0.3771 | 86 | GR | 0.4210 | 933 | 0.4546 |
| 116 | TL | 0.3483 | 772 | 0.3761 | 10 | GS | 0.7706 | 1708 | 0.8321 |
| 117 | DT | 0.3479 | 771 | 0.3756 | 75 | GT | 0.4345 | 963 | 0.4692 |
| 118 | VT | 0.3438 | 762 | 0.3712 | 113 | GV | 0.3564 | 790 | 0.3849 |
| 119 | LR | 0.3415 | 757 | 0.3688 | 329 | GW | 0.0523 | 116 | – |
| 120 | QL | 0.3366 | 746 | 0.3635 | 177 | GY | 0.2224 | 493 | 0.2402 |
| 121 | VK | 0.3352 | 743 | 0.3620 | 253 | HA | 0.1223 | 271 | 0.1320 |
| 122 | RL | 0.3307 | 733 | 0.3571 | 381 | HC | 0.0244 | 54 | – |

Continued on next page

**Table S2 (continued)**

| Rank | Dipeptide | Rel. freq. in DisProt [%] | Abs. occurr. in DisProt | Rel. freq. in dataset [%] | Rank | Dipeptide | Rel. freq. in DisProt [%] | Abs. occurr. in DisProt | Rel. freq. in dataset [%] |
|---|---|---|---|---|---|---|---|---|---|
| Sorted by relative frequency (rank) | | | | | Sorted alphabetically by dipeptide name | | | | |
| 123 | TD | 0.3298 | 731 | 0.3561 | 255 | HD | 0.1205 | 267 | 0.1301 |
| 124 | LT | 0.3276 | 726 | 0.3537 | 237 | HE | 0.1372 | 304 | 0.1481 |
| 125 | NG | 0.3276 | 726 | 0.3537 | 324 | HF | 0.0550 | 122 | – |
| 126 | VG | 0.3249 | 720 | 0.3508 | 206 | HG | 0.1705 | 378 | 0.1842 |
| 127 | DV | 0.3244 | 719 | 0.3503 | 270 | HH | 0.1092 | 242 | 0.1179 |
| 128 | DP | 0.3203 | 710 | 0.3459 | 320 | HI | 0.0596 | 132 | – |
| 129 | TV | 0.3154 | 699 | 0.3406 | 256 | HK | 0.1200 | 266 | 0.1296 |
| 130 | VD | 0.3136 | 695 | 0.3386 | 224 | HL | 0.1498 | 332 | 0.1618 |
| 131 | VV | 0.3050 | 676 | 0.3293 | 355 | HM | 0.0379 | 84 | – |
| 132 | VL | 0.3046 | 675 | 0.3289 | 298 | HN | 0.0835 | 185 | – |
| 133 | PD | 0.3041 | 674 | 0.3284 | 222 | HP | 0.1521 | 337 | 0.1642 |
| 134 | IS | 0.3032 | 672 | 0.3274 | 265 | HQ | 0.1132 | 251 | 0.1223 |
| 135 | KQ | 0.2982 | 661 | 0.3220 | 264 | HR | 0.1132 | 251 | 0.1223 |
| 136 | NE | 0.2978 | 660 | 0.3216 | 194 | HS | 0.1949 | 432 | 0.2105 |
| 137 | RT | 0.2924 | 648 | 0.3157 | 274 | HT | 0.1051 | 233 | 0.1135 |
| 138 | LV | 0.2901 | 643 | 0.3133 | 289 | HV | 0.0902 | 200 | – |
| 139 | NK | 0.2861 | 634 | 0.3089 | 388 | HW | 0.0171 | 38 | – |
| 140 | KN | 0.2815 | 624 | 0.3040 | 352 | HY | 0.0438 | 97 | – |
| 141 | NL | 0.2797 | 620 | 0.3021 | 189 | IA | 0.2066 | 458 | 0.2231 |
| 142 | LN | 0.2793 | 619 | 0.3016 | 374 | IC | 0.0289 | 64 | – |
| 143 | NN | 0.2784 | 617 | 0.3006 | 176 | ID | 0.2229 | 494 | 0.2407 |
| 144 | FG | 0.2743 | 608 | 0.2962 | 163 | IE | 0.2441 | 541 | 0.2636 |
| 145 | NP | 0.2743 | 608 | 0.2962 | 305 | IF | 0.0753 | 167 | – |
| 146 | EI | 0.2721 | 603 | 0.2938 | 190 | IG | 0.2057 | 456 | 0.2222 |
| 147 | NA | 0.2707 | 600 | 0.2923 | 308 | IH | 0.0690 | 153 | – |
| 148 | RQ | 0.2685 | 595 | 0.2899 | 234 | II | 0.1403 | 311 | – |
| 149 | SI | 0.2680 | 594 | 0.2894 | 166 | IK | 0.2351 | 521 | 0.2538 |
| 150 | RD | 0.2653 | 588 | 0.2865 | 185 | IL | 0.2130 | 472 | 0.2300 |
| 151 | QR | 0.2648 | 587 | 0.2860 | 317 | IM | 0.0618 | 137 | – |
| 152 | QT | 0.2617 | 580 | 0.2826 | 225 | IN | 0.1489 | 330 | – |
| 153 | AN | 0.2585 | 573 | 0.2792 | 169 | IP | 0.2319 | 514 | 0.2504 |
| 154 | KI | 0.2567 | 569 | 0.2772 | 214 | IQ | 0.1597 | 354 | 0.1725 |
| 155 | DR | 0.2545 | 564 | 0.2748 | 219 | IR | 0.1543 | 342 | 0.1666 |
| 156 | TQ | 0.2531 | 561 | 0.2733 | 134 | IS | 0.3032 | 672 | 0.3274 |
| 157 | DN | 0.2527 | 560 | 0.2728 | 197 | IT | 0.1918 | 425 | 0.2071 |
| 158 | FS | 0.2495 | 553 | 0.2694 | 213 | IV | 0.1611 | 357 | – |
| 159 | SF | 0.2486 | 551 | 0.2684 | 379 | IW | 0.0248 | 55 | – |
| 160 | TR | 0.2482 | 550 | 0.2680 | 316 | IY | 0.0623 | 138 | – |
| 161 | VQ | 0.2463 | 546 | 0.2660 | 34 | KA | 0.5730 | 1270 | 0.6187 |
| 162 | MA | 0.2445 | 542 | 0.2641 | 342 | KC | 0.0487 | 108 | – |
| 163 | IE | 0.2441 | 541 | 0.2636 | 62 | KD | 0.4598 | 1019 | 0.4965 |
| 164 | NT | 0.2432 | 539 | 0.2626 | 20 | KE | 0.6632 | 1470 | 0.7162 |
| 165 | QD | 0.2369 | 525 | 0.2558 | 231 | KF | 0.1417 | 314 | 0.1530 |
| 166 | IK | 0.2351 | 521 | 0.2538 | 94 | KG | 0.4088 | 906 | 0.4414 |
| 167 | ND | 0.2346 | 520 | 0.2533 | 243 | KH | 0.1317 | 292 | 0.1423 |
| 168 | GF | 0.2328 | 516 | 0.2514 | 154 | KI | 0.2567 | 569 | 0.2772 |
| 169 | IP | 0.2319 | 514 | 0.2504 | 6 | KK | 0.8636 | 1914 | 0.9325 |
| 170 | YS | 0.2310 | 512 | 0.2494 | 93 | KL | 0.4115 | 912 | 0.4443 |
| 171 | GI | 0.2306 | 511 | 0.2490 | 266 | KM | 0.1128 | 250 | 0.1218 |

Continued on next page

**Table S2 (continued)**

| Rank | Dipeptide | Rel. freq. in DisProt [%] | Abs. occurr. in DisProt | Rel. freq. in dataset [%] | Rank | Dipeptide | Rel. freq. in DisProt [%] | Abs. occurr. in DisProt | Rel. freq. in dataset [%] |
|---|---|---|---|---|---|---|---|---|---|
| Sorted by relative frequency (rank) | | | | | Sorted alphabetically by dipeptide name | | | | |
| 172 | TN | 0.2288 | 507 | 0.2470 | 140 | KN | 0.2815 | 624 | 0.3040 |
| 173 | NQ | 0.2256 | 500 | 0.2436 | 46 | KP | 0.5089 | 1128 | 0.5496 |
| 174 | PN | 0.2251 | 499 | 0.2431 | 135 | KQ | 0.2982 | 661 | 0.3220 |
| 175 | DI | 0.2229 | 494 | 0.2407 | 69 | KR | 0.4471 | 991 | 0.4828 |
| 176 | ID | 0.2229 | 494 | 0.2407 | 45 | KS | 0.5098 | 1130 | 0.5505 |
| 177 | GY | 0.2224 | 493 | 0.2402 | 91 | KT | 0.4115 | 912 | 0.4443 |
| 178 | VN | 0.2224 | 493 | 0.2402 | 109 | KV | 0.3659 | 811 | 0.3951 |
| 179 | NV | 0.2220 | 492 | 0.2397 | 373 | KW | 0.0298 | 66 | – |
| 180 | SY | 0.2206 | 489 | 0.2382 | 259 | KY | 0.1169 | 259 | 0.1262 |
| 181 | RN | 0.2202 | 488 | 0.2378 | 48 | LA | 0.5035 | 1116 | 0.5437 |
| 182 | MS | 0.2197 | 487 | 0.2373 | 311 | LC | 0.0654 | 145 | – |
| 183 | QN | 0.2193 | 486 | 0.2368 | 95 | LD | 0.4083 | 905 | 0.4409 |
| 184 | RV | 0.2166 | 480 | 0.2339 | 33 | LE | 0.5753 | 1275 | 0.6212 |
| 185 | IL | 0.2130 | 472 | 0.2300 | 199 | LF | 0.1881 | 417 | 0.2032 |
| 186 | VR | 0.2089 | 463 | 0.2256 | 64 | LG | 0.4575 | 1014 | 0.4940 |
| 187 | QV | 0.2080 | 461 | 0.2246 | 250 | LH | 0.1232 | 273 | 0.1330 |
| 188 | DQ | 0.2071 | 459 | 0.2236 | 198 | LI | 0.1909 | 423 | 0.2061 |
| 189 | IA | 0.2066 | 458 | 0.2231 | 88 | LK | 0.4196 | 930 | 0.4531 |
| 190 | IG | 0.2057 | 456 | 0.2222 | 51 | LL | 0.4986 | 1105 | 0.5384 |
| 191 | AI | 0.2008 | 445 | 0.2168 | 279 | LM | 0.1011 | 224 | – |
| 192 | NR | 0.1999 | 443 | 0.2158 | 142 | LN | 0.2793 | 619 | 0.3016 |
| 193 | YG | 0.1990 | 441 | 0.2149 | 43 | LP | 0.5211 | 1155 | 0.5627 |
| 194 | HS | 0.1949 | 432 | 0.2105 | 112 | LQ | 0.3578 | 793 | 0.3864 |
| 195 | PI | 0.1940 | 430 | 0.2095 | 119 | LR | 0.3415 | 757 | 0.3688 |
| 196 | SH | 0.1936 | 429 | 0.2090 | 24 | LS | 0.6235 | 1382 | 0.6733 |
| 197 | IT | 0.1918 | 425 | 0.2071 | 124 | LT | 0.3276 | 726 | 0.3537 |
| 198 | LI | 0.1909 | 423 | 0.2061 | 138 | LV | 0.2901 | 643 | 0.3133 |
| 199 | LF | 0.1881 | 417 | 0.2032 | 358 | LW | 0.0365 | 81 | – |
| 200 | TI | 0.1854 | 411 | 0.2002 | 263 | LY | 0.1142 | 253 | 0.1233 |
| 201 | RI | 0.1850 | 410 | 0.1998 | 162 | MA | 0.2445 | 542 | 0.2641 |
| 202 | FE | 0.1814 | 402 | 0.1959 | 394 | MC | 0.0113 | 25 | – |
| 203 | ME | 0.1782 | 395 | 0.1924 | 226 | MD | 0.1489 | 330 | 0.1608 |
| 204 | DF | 0.1778 | 394 | 0.1920 | 203 | ME | 0.1782 | 395 | 0.1924 |
| 205 | NI | 0.1760 | 390 | 0.1900 | 336 | MF | 0.0505 | 112 | – |
| 206 | HG | 0.1705 | 378 | 0.1842 | 227 | MG | 0.1466 | 325 | 0.1583 |
| 207 | EF | 0.1701 | 377 | 0.1837 | 360 | MH | 0.0361 | 80 | – |
| 208 | PF | 0.1696 | 376 | 0.1832 | 314 | MI | 0.0641 | 142 | – |
| 209 | AF | 0.1696 | 376 | 0.1832 | 233 | MK | 0.1408 | 312 | 0.1520 |
| 210 | VI | 0.1687 | 374 | 0.1822 | 258 | ML | 0.1178 | 261 | – |
| 211 | FL | 0.1678 | 372 | – | 343 | MM | 0.0483 | 107 | – |
| 212 | GH | 0.1674 | 371 | 0.1808 | 288 | MN | 0.0925 | 205 | – |
| 213 | IV | 0.1611 | 357 | – | 246 | MP | 0.1299 | 288 | 0.1403 |
| 214 | IQ | 0.1597 | 354 | 0.1725 | 299 | MQ | 0.0830 | 184 | – |
| 215 | FP | 0.1588 | 352 | 0.1715 | 297 | MR | 0.0839 | 186 | 0.0906 |
| 216 | FT | 0.1557 | 345 | 0.1681 | 182 | MS | 0.2197 | 487 | 0.2373 |
| 217 | DY | 0.1552 | 344 | 0.1676 | 268 | MT | 0.1101 | 244 | 0.1189 |
| 218 | EH | 0.1552 | 344 | 0.1676 | 293 | MV | 0.0884 | 196 | – |
| 219 | IR | 0.1543 | 342 | 0.1666 | 398 | MW | 0.0081 | 18 | – |
| 220 | EY | 0.1539 | 341 | 0.1661 | 384 | MY | 0.0217 | 48 | – |

Continued on next page

**Table S2 (continued)**

| Sorted by relative frequency (rank) | | | | | Sorted alphabetically by dipeptide name | | | | |
|---|---|---|---|---|---|---|---|---|---|
| Rank | Dipeptide | Rel. freq. in DisProt [%] | Abs. occurr. in DisProt | Rel. freq. in dataset [%] | Rank | Dipeptide | Rel. freq. in DisProt [%] | Abs. occurr. in DisProt | Rel. freq. in dataset [%] |
| 221 | QI | 0.1530 | 339 | 0.1652 | 147 | NA | 0.2707 | 600 | 0.2923 |
| 222 | HP | 0.1521 | 337 | 0.1642 | 362 | NC | 0.0343 | 76 | – |
| 223 | EM | 0.1502 | 333 | 0.1622 | 167 | ND | 0.2346 | 520 | 0.2533 |
| 224 | HL | 0.1498 | 332 | 0.1618 | 136 | NE | 0.2978 | 660 | 0.3216 |
| 225 | IN | 0.1489 | 330 | – | 260 | NF | 0.1155 | 256 | – |
| 226 | MD | 0.1489 | 330 | 0.1608 | 125 | NG | 0.3276 | 726 | 0.3537 |
| 227 | MG | 0.1466 | 325 | 0.1583 | 304 | NH | 0.0772 | 171 | – |
| 228 | FK | 0.1462 | 324 | 0.1579 | 205 | NI | 0.1760 | 390 | 0.1900 |
| 229 | FD | 0.1462 | 324 | 0.1579 | 139 | NK | 0.2861 | 634 | 0.3089 |
| 230 | GM | 0.1457 | 323 | 0.1574 | 141 | NL | 0.2797 | 620 | 0.3021 |
| 231 | KF | 0.1417 | 314 | 0.1530 | 295 | NM | 0.0848 | 188 | – |
| 232 | YE | 0.1412 | 313 | 0.1525 | 143 | NN | 0.2784 | 617 | 0.3006 |
| 233 | MK | 0.1408 | 312 | 0.1520 | 145 | NP | 0.2743 | 608 | 0.2962 |
| 234 | II | 0.1403 | 311 | – | 173 | NQ | 0.2256 | 500 | 0.2436 |
| 235 | FA | 0.1394 | 309 | 0.1505 | 192 | NR | 0.1999 | 443 | 0.2158 |
| 236 | AY | 0.1376 | 305 | 0.1486 | 90 | NS | 0.4133 | 916 | 0.4463 |
| 237 | HE | 0.1372 | 304 | 0.1481 | 164 | NT | 0.2432 | 539 | 0.2626 |
| 238 | PH | 0.1367 | 303 | 0.1476 | 179 | NV | 0.2220 | 492 | 0.2397 |
| 239 | SM | 0.1340 | 297 | 0.1447 | 375 | NW | 0.0275 | 61 | – |
| 240 | YD | 0.1336 | 296 | 0.1442 | 285 | NY | 0.0952 | 211 | – |
| 241 | AH | 0.1331 | 295 | 0.1437 | 12 | PA | 0.7508 | 1664 | 0.8107 |
| 242 | YP | 0.1331 | 295 | 0.1437 | 318 | PC | 0.0605 | 134 | 0.0653 |
| 243 | KH | 0.1317 | 292 | 0.1423 | 133 | PD | 0.3041 | 674 | 0.3284 |
| 244 | TF | 0.1317 | 292 | 0.1423 | 30 | PE | 0.5865 | 1300 | 0.6334 |
| 245 | PY | 0.1308 | 290 | 0.1413 | 208 | PF | 0.1696 | 376 | 0.1832 |
| 246 | MP | 0.1299 | 288 | 0.1403 | 27 | PG | 0.6087 | 1349 | 0.6572 |
| 247 | YL | 0.1290 | 286 | 0.1393 | 238 | PH | 0.1367 | 303 | 0.1476 |
| 248 | AM | 0.1263 | 280 | 0.1364 | 195 | PI | 0.1940 | 430 | 0.2095 |
| 249 | FQ | 0.1236 | 274 | 0.1335 | 59 | PK | 0.4724 | 1047 | 0.5101 |
| 250 | LH | 0.1232 | 273 | 0.1330 | 67 | PL | 0.4503 | 998 | 0.4862 |
| 251 | RH | 0.1227 | 272 | 0.1325 | 257 | PM | 0.1182 | 262 | 0.1276 |
| 252 | DH | 0.1223 | 271 | 0.1320 | 174 | PN | 0.2251 | 499 | 0.2431 |
| 253 | HA | 0.1223 | 271 | 0.1320 | 7 | PP | 0.8464 | 1876 | 0.9140 |
| 254 | VF | 0.1209 | 268 | 0.1306 | 87 | PQ | 0.4210 | 933 | 0.4546 |
| 255 | HD | 0.1205 | 267 | 0.1301 | 105 | PR | 0.3713 | 823 | 0.4010 |
| 256 | HK | 0.1200 | 266 | 0.1296 | 8 | PS | 0.7995 | 1772 | 0.8633 |
| 257 | PM | 0.1182 | 262 | 0.1276 | 47 | PT | 0.5076 | 1125 | 0.5481 |
| 258 | ML | 0.1178 | 261 | – | 71 | PV | 0.4467 | 990 | 0.4823 |
| 259 | KY | 0.1169 | 259 | 0.1262 | 339 | PW | 0.0492 | 109 | 0.0531 |
| 260 | NF | 0.1155 | 256 | – | 245 | PY | 0.1308 | 290 | 0.1413 |
| 261 | YN | 0.1151 | 255 | – | 98 | QA | 0.3912 | 867 | 0.4224 |
| 262 | QH | 0.1142 | 253 | 0.1233 | 366 | QC | 0.0316 | 70 | – |
| 263 | LY | 0.1142 | 253 | 0.1233 | 165 | QD | 0.2369 | 525 | 0.2558 |
| 264 | HR | 0.1132 | 251 | 0.1223 | 79 | QE | 0.4340 | 962 | 0.4687 |
| 265 | HQ | 0.1132 | 251 | 0.1223 | 286 | QF | 0.0943 | 209 | 0.1018 |
| 266 | KM | 0.1128 | 250 | 0.1218 | 111 | QG | 0.3582 | 794 | 0.3868 |
| 267 | YT | 0.1119 | 248 | – | 262 | QH | 0.1142 | 253 | 0.1233 |
| 268 | MT | 0.1101 | 244 | 0.1189 | 221 | QI | 0.1530 | 339 | 0.1652 |
| 269 | FV | 0.1101 | 244 | – | 115 | QK | 0.3492 | 774 | 0.3771 |

Continued on next page

**Table S2 (continued)**

| Sorted by relative frequency (rank) | | | | | Sorted alphabetically by dipeptide name | | | | |
|---|---|---|---|---|---|---|---|---|---|
| Rank | Dipep-tide | Rel. freq. in DisProt [%] | Abs. occurr. in DisProt | Rel. freq. in dataset [%] | Rank | Dipep-tide | Rel. freq. in DisProt [%] | Abs. occurr. in DisProt | Rel. freq. in dataset [%] |
| 270 | HH | 0.1092 | 242 | 0.1179 | 120 | QL | 0.3366 | 746 | 0.3635 |
| 271 | FR | 0.1087 | 241 | 0.1174 | 300 | QM | 0.0821 | 182 | – |
| 272 | TY | 0.1065 | 236 | 0.1150 | 183 | QN | 0.2193 | 486 | 0.2368 |
| 273 | FN | 0.1056 | 234 | – | 77 | QP | 0.4345 | 963 | 0.4692 |
| 274 | HT | 0.1051 | 233 | 0.1135 | 53 | QQ | 0.4846 | 1074 | 0.5233 |
| 275 | YR | 0.1047 | 232 | 0.1130 | 151 | QR | 0.2648 | 587 | 0.2860 |
| 276 | TH | 0.1042 | 231 | 0.1125 | 96 | QS | 0.3961 | 878 | 0.4278 |
| 277 | DM | 0.1024 | 227 | – | 152 | QT | 0.2617 | 580 | 0.2826 |
| 278 | RF | 0.1020 | 226 | – | 187 | QV | 0.2080 | 461 | 0.2246 |
| 279 | LM | 0.1011 | 224 | – | 368 | QW | 0.0307 | 68 | – |
| 280 | YQ | 0.0997 | 221 | 0.1077 | 291 | QY | 0.0889 | 197 | – |
| 281 | YK | 0.0993 | 220 | 0.1072 | 114 | RA | 0.3546 | 786 | 0.3829 |
| 282 | CS | 0.0993 | 220 | – | 354 | RC | 0.0433 | 96 | – |
| 283 | YA | 0.0988 | 219 | – | 150 | RD | 0.2653 | 588 | 0.2865 |
| 284 | VH | 0.0970 | 215 | 0.1047 | 97 | RE | 0.3948 | 875 | 0.4263 |
| 285 | NY | 0.0952 | 211 | – | 278 | RF | 0.1020 | 226 | – |
| 286 | QF | 0.0943 | 209 | 0.1018 | 82 | RG | 0.4277 | 948 | 0.4619 |
| 287 | TM | 0.0938 | 208 | 0.1013 | 251 | RH | 0.1227 | 272 | 0.1325 |
| 288 | MN | 0.0925 | 205 | – | 201 | RI | 0.1850 | 410 | 0.1998 |
| 289 | HV | 0.0902 | 200 | – | 83 | RK | 0.4264 | 945 | 0.4604 |
| 290 | YV | 0.0889 | 197 | – | 122 | RL | 0.3307 | 733 | 0.3571 |
| 291 | QY | 0.0889 | 197 | – | 296 | RM | 0.0839 | 186 | – |
| 292 | SC | 0.0884 | 196 | – | 181 | RN | 0.2202 | 488 | 0.2378 |
| 293 | MV | 0.0884 | 196 | – | 107 | RP | 0.3673 | 814 | 0.3966 |
| 294 | RY | 0.0880 | 195 | – | 148 | RQ | 0.2685 | 595 | 0.2899 |
| 295 | NM | 0.0848 | 188 | – | 44 | RR | 0.5175 | 1147 | 0.5588 |
| 296 | RM | 0.0839 | 186 | – | 50 | RS | 0.4986 | 1105 | 0.5384 |
| 297 | MR | 0.0839 | 186 | 0.0906 | 137 | RT | 0.2924 | 648 | 0.3157 |
| 298 | HN | 0.0835 | 185 | – | 184 | RV | 0.2166 | 480 | 0.2339 |
| 299 | MQ | 0.0830 | 184 | – | 356 | RW | 0.0370 | 82 | – |
| 300 | QM | 0.0821 | 182 | – | 294 | RY | 0.0880 | 195 | – |
| 301 | VY | 0.0812 | 180 | – | 14 | SA | 0.7178 | 1591 | 0.7751 |
| 302 | FI | 0.0794 | 176 | – | 292 | SC | 0.0884 | 196 | – |
| 303 | FF | 0.0790 | 175 | – | 31 | SD | 0.5843 | 1295 | 0.6309 |
| 304 | NH | 0.0772 | 171 | – | 17 | SE | 0.6786 | 1504 | 0.7328 |
| 305 | IF | 0.0753 | 167 | – | 159 | SF | 0.2486 | 551 | 0.2684 |
| 306 | CL | 0.0735 | 163 | – | 9 | SG | 0.7927 | 1757 | 0.8560 |
| 307 | VM | 0.0708 | 157 | – | 196 | SH | 0.1936 | 429 | 0.2090 |
| 308 | IH | 0.0690 | 153 | – | 149 | SI | 0.2680 | 594 | 0.2894 |
| 309 | CG | 0.0663 | 147 | – | 36 | SK | 0.5708 | 1265 | 0.6163 |
| 310 | CP | 0.0659 | 146 | 0.0711 | 28 | SL | 0.6055 | 1342 | 0.6538 |
| 311 | LC | 0.0654 | 145 | – | 239 | SM | 0.1340 | 297 | 0.1447 |
| 312 | YI | 0.0654 | 145 | – | 110 | SN | 0.3659 | 811 | 0.3951 |
| 313 | YY | 0.0641 | 142 | – | 5 | SP | 0.8816 | 1954 | 0.9520 |
| 314 | MI | 0.0641 | 142 | – | 68 | SQ | 0.4489 | 995 | 0.4848 |
| 315 | GC | 0.0627 | 139 | – | 40 | SR | 0.5369 | 1190 | 0.5798 |
| 316 | IY | 0.0623 | 138 | – | 2 | SS | 1.3224 | 2931 | 1.4280 |
| 317 | IM | 0.0618 | 137 | – | 32 | ST | 0.5811 | 1288 | 0.6275 |
| 318 | PC | 0.0605 | 134 | 0.0653 | 84 | SV | 0.4232 | 938 | 0.4570 |

Continued on next page

**Table S2 (continued)**

| Sorted by relative frequency (rank) | | | | | Sorted alphabetically by dipeptide name | | | | |
|---|---|---|---|---|---|---|---|---|---|
| Rank | Dipep-tide | Rel. freq. in DisProt [%] | Abs. occurr. in DisProt | Rel. freq. in dataset [%] | Rank | Dipep-tide | Rel. freq. in DisProt [%] | Abs. occurr. in DisProt | Rel. freq. in dataset [%] |
| 319 | WG | 0.0600 | 133 | – | 326 | SW | 0.0528 | 117 | – |
| 320 | HI | 0.0596 | 132 | – | 180 | SY | 0.2206 | 489 | 0.2382 |
| 321 | AW | 0.0591 | 131 | – | 49 | TA | 0.5004 | 1109 | 0.5403 |
| 322 | EC | 0.0591 | 131 | – | 335 | TC | 0.0510 | 113 | – |
| 323 | AC | 0.0559 | 124 | – | 123 | TD | 0.3298 | 731 | 0.3561 |
| 324 | HF | 0.0550 | 122 | – | 60 | TE | 0.4715 | 1045 | 0.5091 |
| 325 | YF | 0.0541 | 120 | – | 244 | TF | 0.1317 | 292 | 0.1423 |
| 326 | SW | 0.0528 | 117 | – | 80 | TG | 0.4327 | 959 | 0.4672 |
| 327 | FH | 0.0528 | 117 | – | 276 | TH | 0.1042 | 231 | 0.1125 |
| 328 | CT | 0.0523 | 116 | – | 200 | TI | 0.1854 | 411 | 0.2002 |
| 329 | GW | 0.0523 | 116 | – | 100 | TK | 0.3853 | 854 | 0.4161 |
| 330 | WE | 0.0519 | 115 | – | 116 | TL | 0.3483 | 772 | 0.3761 |
| 331 | CD | 0.0514 | 114 | – | 287 | TM | 0.0938 | 208 | 0.1013 |
| 332 | VC | 0.0510 | 113 | – | 172 | TN | 0.2288 | 507 | 0.2470 |
| 333 | CK | 0.0510 | 113 | – | 38 | TP | 0.5473 | 1213 | 0.5910 |
| 334 | FY | 0.0510 | 113 | – | 156 | TQ | 0.2531 | 561 | 0.2733 |
| 335 | TC | 0.0510 | 113 | – | 160 | TR | 0.2482 | 550 | 0.2680 |
| 336 | MF | 0.0505 | 112 | – | 19 | TS | 0.6687 | 1482 | 0.7220 |
| 337 | YH | 0.0496 | 110 | – | 73 | TT | 0.4408 | 977 | 0.4760 |
| 338 | WD | 0.0496 | 110 | – | 129 | TV | 0.3154 | 699 | 0.3406 |
| 339 | PW | 0.0492 | 109 | 0.0531 | 365 | TW | 0.0325 | 72 | – |
| 340 | WA | 0.0492 | 109 | – | 272 | TY | 0.1065 | 236 | 0.1150 |
| 341 | DW | 0.0487 | 108 | – | 92 | VA | 0.4115 | 912 | 0.4443 |
| 342 | KC | 0.0487 | 108 | – | 332 | VC | 0.0510 | 113 | – |
| 343 | MM | 0.0483 | 107 | – | 130 | VD | 0.3136 | 695 | 0.3386 |
| 344 | CE | 0.0474 | 105 | – | 78 | VE | 0.4340 | 962 | 0.4687 |
| 345 | WS | 0.0469 | 104 | – | 254 | VF | 0.1209 | 268 | 0.1306 |
| 346 | CR | 0.0465 | 103 | – | 126 | VG | 0.3249 | 720 | 0.3508 |
| 347 | EW | 0.0465 | 103 | – | 284 | VH | 0.0970 | 215 | 0.1047 |
| 348 | WL | 0.0456 | 101 | – | 210 | VI | 0.1687 | 374 | 0.1822 |
| 349 | DC | 0.0456 | 101 | – | 121 | VK | 0.3352 | 743 | 0.3620 |
| 350 | CA | 0.0456 | 101 | – | 132 | VL | 0.3046 | 675 | 0.3289 |
| 351 | CV | 0.0442 | 98 | – | 307 | VM | 0.0708 | 157 | – |
| 352 | HY | 0.0438 | 97 | – | 178 | VN | 0.2224 | 493 | 0.2402 |
| 353 | WK | 0.0433 | 96 | – | 70 | VP | 0.4467 | 990 | 0.4823 |
| 354 | RC | 0.0433 | 96 | – | 161 | VQ | 0.2463 | 546 | 0.2660 |
| 355 | HM | 0.0379 | 84 | – | 186 | VR | 0.2089 | 463 | 0.2256 |
| 356 | RW | 0.0370 | 82 | – | 65 | VS | 0.4521 | 1002 | 0.4882 |
| 357 | WR | 0.0370 | 82 | – | 118 | VT | 0.3438 | 762 | 0.3712 |
| 358 | LW | 0.0365 | 81 | – | 131 | VV | 0.3050 | 676 | 0.3293 |
| 359 | WN | 0.0365 | 81 | – | 371 | VW | 0.0302 | 67 | – |
| 360 | MH | 0.0361 | 80 | – | 301 | VY | 0.0812 | 180 | – |
| 361 | FM | 0.0356 | 79 | – | 340 | WA | 0.0492 | 109 | – |
| 362 | NC | 0.0343 | 76 | – | 399 | WC | 0.0072 | 16 | – |
| 363 | CQ | 0.0334 | 74 | – | 338 | WD | 0.0496 | 110 | – |
| 364 | WT | 0.0325 | 72 | – | 330 | WE | 0.0519 | 115 | – |
| 365 | TW | 0.0325 | 72 | – | 386 | WF | 0.0208 | 46 | – |
| 366 | QC | 0.0316 | 70 | – | 319 | WG | 0.0600 | 133 | – |
| 367 | WQ | 0.0316 | 70 | – | 393 | WH | 0.0131 | 29 | – |

Continued on next page

**Table S2 (continued)**

| Sorted by relative frequency (rank) | | | | | Sorted alphabetically by dipeptide name | | | | |
|---|---|---|---|---|---|---|---|---|---|
| Rank | Dipep-tide | Rel. freq. in DisProt [%] | Abs. occurr. in DisProt | Rel. freq. in dataset [%] | Rank | Dipep-tide | Rel. freq. in DisProt [%] | Abs. occurr. in DisProt | Rel. freq. in dataset [%] |
| 368 | QW | 0.0307 | 68 | – | 389 | WI | 0.0167 | 37 | – |
| 369 | CN | 0.0307 | 68 | – | 353 | WK | 0.0433 | 96 | – |
| 370 | YM | 0.0302 | 67 | – | 348 | WL | 0.0456 | 101 | – |
| 371 | VW | 0.0302 | 67 | – | 395 | WM | 0.0104 | 23 | – |
| 372 | CI | 0.0298 | 66 | – | 359 | WN | 0.0365 | 81 | – |
| 373 | KW | 0.0298 | 66 | – | 377 | WP | 0.0257 | 57 | 0.0278 |
| 374 | IC | 0.0289 | 64 | – | 367 | WQ | 0.0316 | 70 | – |
| 375 | NW | 0.0275 | 61 | – | 357 | WR | 0.0370 | 82 | – |
| 376 | FC | 0.0266 | 59 | – | 345 | WS | 0.0469 | 104 | – |
| 377 | WP | 0.0257 | 57 | 0.0278 | 364 | WT | 0.0325 | 72 | – |
| 378 | YC | 0.0257 | 57 | – | 380 | WV | 0.0248 | 55 | – |
| 379 | IW | 0.0248 | 55 | – | 396 | WW | 0.0099 | 22 | – |
| 380 | WV | 0.0248 | 55 | – | 392 | WY | 0.0135 | 30 | – |
| 381 | HC | 0.0244 | 54 | – | 283 | YA | 0.0988 | 219 | – |
| 382 | CF | 0.0235 | 52 | – | 378 | YC | 0.0257 | 57 | – |
| 383 | CC | 0.0230 | 51 | – | 240 | YD | 0.1336 | 296 | 0.1442 |
| 384 | MY | 0.0217 | 48 | – | 232 | YE | 0.1412 | 313 | 0.1525 |
| 385 | CY | 0.0217 | 48 | – | 325 | YF | 0.0541 | 120 | – |
| 386 | WF | 0.0208 | 46 | – | 193 | YG | 0.1990 | 441 | 0.2149 |
| 387 | CH | 0.0180 | 40 | – | 337 | YH | 0.0496 | 110 | – |
| 388 | HW | 0.0171 | 38 | – | 312 | YI | 0.0654 | 145 | – |
| 389 | WI | 0.0167 | 37 | – | 281 | YK | 0.0993 | 220 | 0.1072 |
| 390 | CM | 0.0162 | 36 | – | 247 | YL | 0.1290 | 286 | 0.1393 |
| 391 | FW | 0.0149 | 33 | – | 370 | YM | 0.0302 | 67 | – |
| 392 | WY | 0.0135 | 30 | – | 261 | YN | 0.1151 | 255 | – |
| 393 | WH | 0.0131 | 29 | – | 242 | YP | 0.1331 | 295 | 0.1437 |
| 394 | MC | 0.0113 | 25 | – | 280 | YQ | 0.0997 | 221 | 0.1077 |
| 395 | WM | 0.0104 | 23 | – | 275 | YR | 0.1047 | 232 | 0.1130 |
| 396 | WW | 0.0099 | 22 | – | 170 | YS | 0.2310 | 512 | 0.2494 |
| 397 | YW | 0.0099 | 22 | – | 267 | YT | 0.1119 | 248 | – |
| 398 | MW | 0.0081 | 18 | – | 290 | YV | 0.0889 | 197 | – |
| 399 | WC | 0.0072 | 16 | – | 397 | YW | 0.0099 | 22 | – |
| 400 | CW | 0.0054 | 12 | – | 313 | YY | 0.0641 | 142 | – |

**Table S3.** Average chemical shift differences between protonated (pH $\approx$ 3) and deprotonated (pH $\approx$ 6) ASP and GLU residues. The protonation-dependent absolute differences were calculated separately as $\Delta\delta = |\delta_{\text{prot}} - \delta_{\text{deprot}}|$ for $^{1}$H, $^{13}$C, and $^{15}$N, and averaged over all corresponding nuclei (n) in dipeptides containing ASP or GLU. All reported differences exceed the typical assignment uncertainties of the respective nuclei ($^{1}$H: 0.01 ppm, $^{13}$C: 0.05 ppm, and $^{15}$N: 0.1 ppm).

| | **$^{1}$H** | | **$^{13}$C** | | **$^{15}$N** | |
|---|---|---|---|---|---|---|
| **Average over** | $\Delta\delta$ [ppm] | **n** | $\Delta\delta$ [ppm] | **n** | $\Delta\delta$ [ppm] | **n** |
| ASP | 0.2544 ± 0.1224 | 140 | 2.7416 ± 1.0128 | 140 | 3.5798 ± 2.3160 | 35 |
| GLU | 0.1128 ± 0.0669 | 216 | 2.0175 ± 1.3897 | 180 | 0.6842 ± 0.2202 | 36 |
| ASP and GLU | 0.1685 ± 0.1157 | 356 | 2.3343 ± 1.2900 | 320 | 2.1116 ± 2.1827 | 71 |

**Table S4.** Average chemical shift differences determined for the partner (non-GLU/ASP) residues in dipeptides containing ASP or GLU. Differences measured between dipeptides with protonated (pH $\approx$ 3) and deprotonated (pH $\approx$ 6) ASP or GLU side-chain carboxyl groups. Absolute differences were calculated for $^{1}$H, $^{13}$C, and $^{15}$N as $\Delta\delta = |\delta_{\text{prot}} - \delta_{\text{deprot}}|$ and averaged over all corresponding nuclei (n). All reported differences exceed the typical assignment uncertainties of the respective nuclei ($^{1}$H: 0.01 ppm, $^{13}$C: 0.05 ppm, and $^{15}$N: 0.1 ppm).

| | **$^{1}$H** | | **$^{13}$C** | | **$^{15}$N** | |
|---|---|---|---|---|---|---|
| **Average over** | $\Delta\delta$ [ppm] | **n** | $\Delta\delta$ [ppm] | **n** | $\Delta\delta$ [ppm] | **n** |
| ASP | 0.0346 ± 0.0468 | 292 | 0.2482 ± 0.3370 | 225 | 0.2228 ± 0.1789 | 64 |
| GLU | 0.0138 ± 0.0213 | 302 | 0.0780 ± 0.1489 | 232 | 0.1278 ± 0.0972 | 65 |
| ASP and GLU | 0.0240 ± 0.0376 | 594 | 0.1618 ± 0.2728 | 457 | 0.1753 ± 0.1516 | 129 |

### References


**1.** Aspromonte, M. C. *et al.* DisProt in 2024: Improving function annotation of intrinsically disordered proteins. *Nucleic Acids Res.* **52**, D434–D441 (2024).

**2.** Hoch, J. C. *et al.* Biological magnetic resonance data bank. *Nucleic Acids Res.* **51**, D368–D376 (2023).
